# Specificity for deubiquitination of monoubiquitinated FANCD2 is driven by the N-terminus of USP1

**DOI:** 10.1101/430124

**Authors:** Connor Arkinson, Viduth K. Chaugule, Rachel Toth, Helen Walden

## Abstract

The DNA damage response depends on ubiquitin signalling to orchestrate DNA repair. The Fanconi Anemia pathway for interstrand crosslink repair, and the translesion synthesis pathway for DNA damage tolerance, both require cycles of monoubiquitination and deubiquitination. The ubiquitin specific protease USP1 regulates both these pathways by deubiquitinating monoubiquitinated PCNA, FANCD2 and FANCI. Loss of USP1 activity gives rise to chromosomal instability. While many USPs hydrolyse ubiquitin-ubiquitin linkages, USP1 targets ubiquitin-substrate conjugates at specific sites. The molecular basis of USP1’s specificity for multiple substrates is poorly understood. Here we show that the molecular determinants for substrate deubiquitination by USP1 reside within the highly conserved and extended N-terminus. We find that the N-terminus of USP1 harbours a FANCD2-specific binding sequence required for deubiquitination of K561 on FANCD2. In contrast, the N-terminus is not required for PCNA or FANCI deubiquitination. Furthermore, we show that the N-terminus of USP1 is sufficient to engineer specificity in a more promiscuous USP.

## Introduction

Ubiquitination is a reversible post-translational modification that regulates almost every cellular process in eukaryotes. Cycles of ubiquitination and deubiquitination orchestrate the assembly and disassembly of many DNA repair complexes in DNA damage response pathways. These include the Fanconi Anemia (FA) pathway, required to repair DNA interstrand crosslinks (ICLs), and the translesion synthesis pathway (TLS), required for DNA damage tolerance^1^. FA is a chromosomal instability disorder that results from a dysfunctional ICL repair pathway^2^. Central to FA ICL repair is monoubiquitination of two homologous proteins, FANCD2 and FANCI, at two specific lysines, K561 and K523, respectively, catalysed by FANCL E3 and Ube2T E2^3^. Monoubiquitinated FANCD2 (FANCD2-Ub) signals multiple DNA repair proteins to conduct ICL repair^2^. A similar specific modification is central to TLS repair, where K164 of PCNA is monoubiquitinated (PCNA-Ub) by the RAD18 E3 ligase and Rad6 E2^4^. PCNA-Ub recruits TLS polymerases for DNA repair^5^. As well as ubiquitination, both ICL and TLS repair require deubiquitination (removal of ubiquitin). Interestingly, although the modifications in each pathway are assembled by distinct enzymes, they are removed by the same deubiquitinase, the USP1-UAF1 complex^6^-^8^8. Loss of USP1 function results in an accumulation of monoubiquitinated FANCD2, FANCI and PCNA, genomic instability and a failure to complete the pathways^7,9^-12. In addition to these three substrates, USP1 deubiquitinates a number of other substrates including inhibitor of DNA binding proteins 1–4 (ID1–4) which regulate cell differentiation^13^ and TBK1 which is involved in viral infection^14^.

USP1 belongs to the largest family of deubiquitinases (DUBs), the ubiquitin specific proteases (USPs), which contain ∼50 members. Many USPs show little substrate discrimination between ubiquitin-ubiquitin chains *in vitro^15^*, and can hydrolyse polyubiquitin chains from substrates^16^. A few USPs exhibit preference for specific ubiquitin-ubiquitin linkages such as USP30 for K6-linked Ub chains^17^. In contrast, USP1 targets monoubiquitinated substrates, and regulates a distinct set of modified proteins. Although molecular mechanisms of ubiquitin removal from ubiquitin are well understood, it is less clear how ubiquitin-substrate linkages are specifically targeted. The core catalytic USP domain is ∼350 amino acids. However, most USPs also contain multiple insertions within the catalytic domain and additional N/C- terminal extensions^18^. USP1 has multiple insertions and an extended N-terminus on its USP domain, their functions are currently unknown.

USP1 has little DUB activity on its own, but is regulated by and forms a stoichiometric complex with UAF1. UAF1 also binds and activates two other DUBs, USP12 and USP46^19^, and studies that reveal how UAF1 binds and activates USP12 and USP46 suggest that UAF1 will bind to USP1 in an analogous manner^20,21^. UAF1 acts to stabilise its USP partners and increase catalytic activity^22^, and UAF1 knockout in mice is embryonic lethal, while USP1 knockouts result in a FA-like phenotype, reflecting the additional functions of UAF1^9,23^. In addition to its activation role, UAF1 has a C-terminal Sumo-like domain (SLD) responsible for recruiting USP1 indirectly to FANCD2 and PCNA via a weak interaction with FANCI and ATAD5, respectively^24^. Despite the common activator function of UAF1, loss of either USP12 or USP46 does not result in accumulation of USP1 substrates^19^, thus it remains unclear how USP1 specifically targets its substrate pool.

Investigating how USPs deubiquitinate their substrates on a molecular level is very challenging due to the difficulty in making physiological and correctly ubiquitinated substrates. To date most of our understanding of DUB specificity has used ubiquitin-ubiquitin linkages as substrates, likely due to the advances in purifying large quantities of ubiquitin chains. However, there are few examples of studies which have used monoubiquitinated substrates with a native isopeptide, and these include PCNA-Ub^25^ and histones^26^. Particularly in the case of histone H2A and H2B, the generation of monoubiquitinated substrates has facilitated the elucidation of the mechanisms of substrate targeting by histone-specific DUBs^26,27^. The ability to make physiological substrates allows for a modular approach to understanding the requirements for specificity and how DUBs such as USP1 work at the molecular level.

Here we report the reconstitution of substrate deubiquitination by USP1-UAF1 that allows for a modular approach to understanding the molecular requirements for deubiquitination of physiological substrates. We define the molecular determinants for substrate deubiquitination via a few residues in a highly conserved and extended N-terminus of USP1. Our analysis indicates that the N-terminus of USP1 harbours a FANCD2 specific binding sequence important for only deubiquitinating one specific lysine on FANCD2, K561 – the location of a specific DNA repair signal. Remarkably, we find that the N-terminus of USP1 is important for FANCD2-K561Ub deubiquitination but not for PCNA-K164Ub or FANCI-K523Ub. Finally, we find that the N-terminus of USP1 is sufficient to engineer specificity in a more promiscuous DUB. Our analysis show that USP1 discriminates between substrates and has direct elements for targeting FANCD2, regardless of whether it is ubiquitinated providing further insights into how USPs select their substrates.

## Results

### USP1 catalytic domain is sufficient for activity and binding to UAF1

To gain insight into USP1 specificity, we analysed the amino acid sequence and separated the protein into regions termed USP domain, Insert 1, Insert 2 and N-terminus (Figure 1A). Several elements important for cellular regulation have been previously identified; such as a calpain cleavage site within the N-terminus^28^, a degron motif for APC/C targeting within insertion 1^29^ and an auto-cleavage region (G670/G671) within insertion 2^7^. However, whether these regions are important for *in vitro* DUB activity is unknown. We designed multiple USP1 fragments for expression in sf21 cells, keeping the catalytic domain intact, but systematically removing the insertions and N-terminus (Figure 1B). Each construct of USP1 is expressed and purified to homogeneity (Figure 1C). In order to assess catalytic activity and competency we used Ubiquitin-propargylamine (Ub-prg), a suicide probe which crosslinks to the active site cysteine residue of DUBs^30^. Recombinant USP1^FL^ in addition to all of the other fragments fully reacts with Ub-prg (Figure 1C), thus indicating the catalytic domain is competent and able to bind ubiquitin. USP1^FL^ has multiple break down products, but when insertion 1 and 2 are deleted, the yield and purity is much higher, indicating an increase of protein stability. Indeed, thermal denaturation assays reveal that USP1^ΔNΔ1Δ2^ is more thermostable (Tm = 44 ± 0.24°C, whereas the USP1^FL^ melts at lower temperatures (Tm = 37 ± 0.81°C (Figure EV1). Importantly, USP1 requires to be in complex with UAF1 for robust catalytic activity^22^. We find that the minimal USP1 catalytic domain (USP1^ΔNΔ1Δ2^) is sufficient for binding to UAF1 by size exclusion chromatography (Figure 1D) and stimulating activity of USP1 on a fluorescent ubiquitinated-dipeptide substrate, Ub-KG^TAMRA^ (Figure 1E). The cleavage rates shows that the activity of USP1^ΔNΔ1Δ2^ is almost identical to USP1^FL^-UAF1. Since a minimal pseudo substrate was used, we further assayed the activity against K63 and K48 linked di-ubiquitin chains (Figure 1F) and do not detect any differences in activity between USP1^FL^ and USP1^ΔNΔ1Δ2^. Together, these data show that the catalytic domain of USP1, with stoichiometric amounts of UAF1, is sufficient for DUB activity, and that the additional regions of USP1 are not important for *in vitro* DUB catalysis or UAF1 stimulated activity.

**Figure 1.**
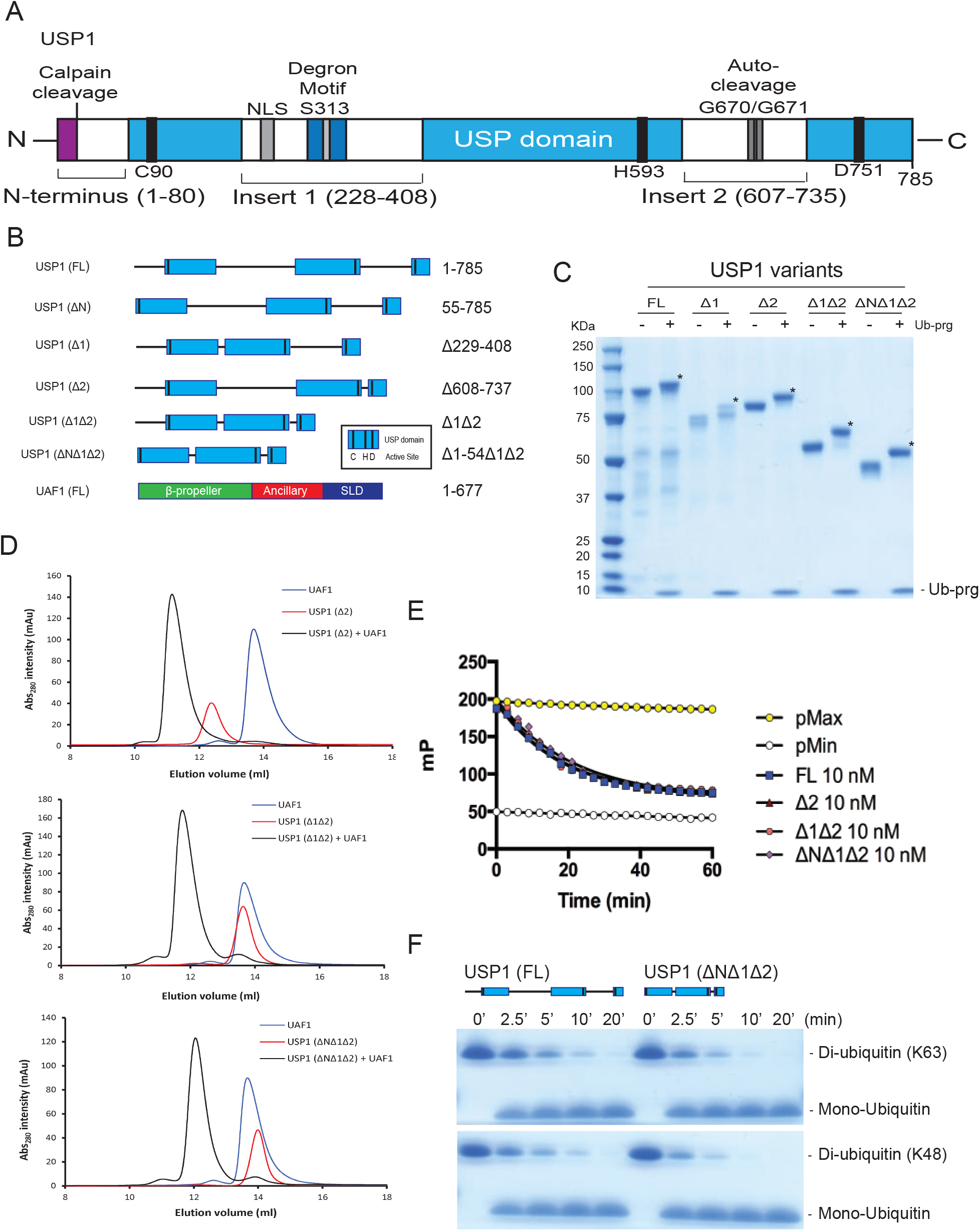
The catalytic domain of USP1 is sufficent for UAF1 binding and activation. A) Overall schematic of USP1 domain structure and boundaries. USP1 contains an extended N-terminus and two large insertions within the USP domain. This schematic excludes minor insertions within the catalytic domain. NLS; nuclear localisation signal B) Scehmatic of truncation/deletion constructs of USP1 fragments and UAF1 used in this study. C) Recombinant USP1 constructs from sf21 cells incubated with 3-fold molar excess Ubiquitin-propargylated (ub-prg). A band shift (*) shows crosslinking between USP1 and Ubiquitin-prg. D) Size exclusion chromatography of recombinant USP1 with UAF1 at 15 µM. Catalytic domain of USP1 (USP1^ΔNΔ1Δ2^) is sufficent for complex with UAF1. E) A comparison of deubiquitinase activity of USP1 constructs using [300 nM] Ub-KG-TAMRA as a minimal substrate. Equimolar UAF1 was used as USP1 alone contains little activity. F) Hydrolysis of K63 or K48 linked di-ubiquitin (10 µM) using 50 nM USP1^FL^-UAF1 or 50 nM USP1^ΔN^Δ1Δ2-UAF1. Data shows comparable activity of FL USP1 and Catalytic domain of USP1. FL = full length.

**Figure EV1.**
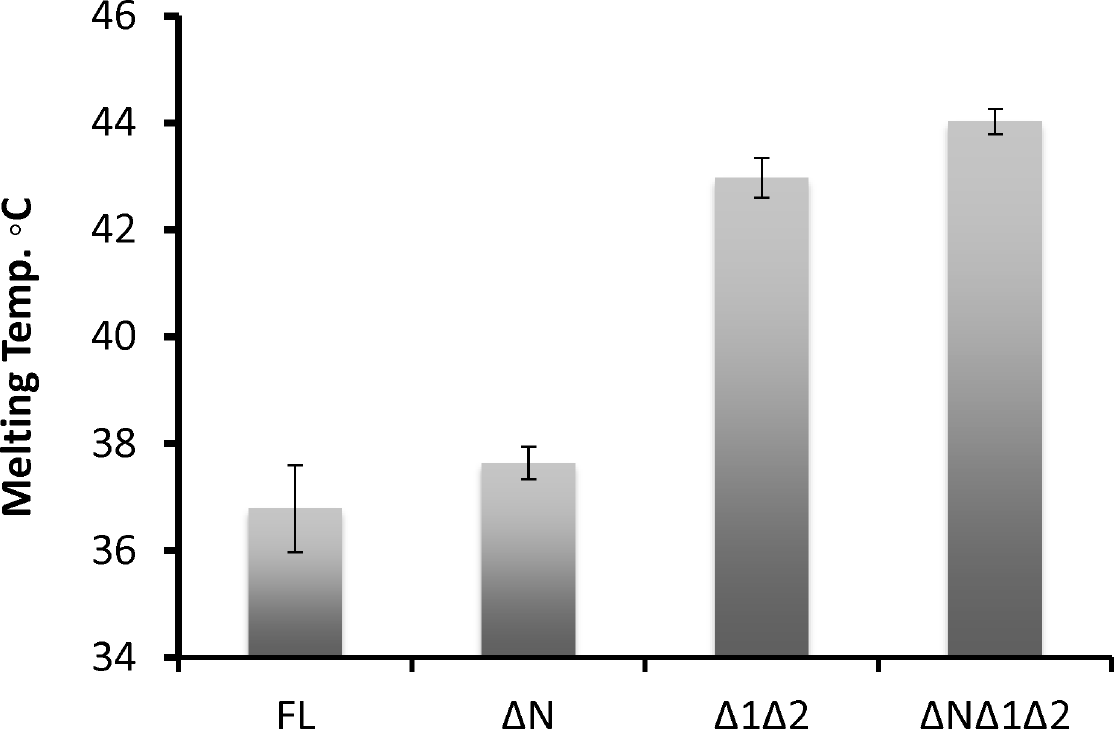
Thermo shift denaturation assay of recombinant USP1 variants. Data shows mean melting temperature from [n = 9] +/− SD of USP1 variants using sypro-organge dye. USP1^FL^, USP1^ΔN^ (Δ1–54), USP1^Δ1Δ2^ (Δ229–408, Δ608–737) and USP1^ΔNΔ1Δ2^ (Δ1–54, Δ229–408, Δ608–737).

### USP1-UAF1 directly deubiquitinates monomeric FANCD2-Ub and FANCI-Ub

The USP1-UAF1 complex can deubiquitinate several monoubiquitinated substrates, including human FANCD2-Ub (*hs*FANCD2-Ub)^6^, human FANCI-Ub (*hs*FANCI-Ub)^31^ and human (*hs*PCNA-Ub)^25^. In order to understand the requirements for deubiquitination we optimised a robust method for producing full length isolated *hs*FANCD2-Ub and *hs*FANCI-Ub substrates. The homogenous preparation of isolated *hs*FANCD2-Ub and *hs*FANCI-Ub is very challenging, as studies required non-mammalian FANCD2 and FANCI complexes, in addition to needing DNA, and a 6 protein complex (FANCC, FANCE, FANCF, FANCB, FANCL and FAAP100) to stimulate the modification^32^. To enhance the yield and purity of *hs*FANCD2-Ub and *hs*FANCI-Ub, we developed a method that requires a FANCL fragment (N-terminally truncated), a hyperactive mutant form of Ube2T^V3^33, E1 and ubiquitin. Using these enzymes (Figure 2A), we observe robust monoubiquitination of the isolated substrates *hs*FANCD2 and *hs*FANCI substrates within 60 min when monitored with either fluorescent ubiquitin (Ub^800^) (Figure EV2) or GST-Ub (Figure 2B). In order to ensure site-specificity of ubiquitination, we mutated K561 and K523 to Arg in *hs*FANCD2 and *hs*FANCI, respectively. In contrast to wild-type (WT) *hs*FANCD2 and *hs*FANCI, K561R and K523R is not monoubiquitinated using these reaction conditions, indicating site specificity is maintained (Figure 2B). We further assessed site-specificity by purifying *hs*FANCD2-Ub and *hs*FANCI-Ub to homogeneity and analysed both modified and un-modified proteins by mass spectrometry to verify is the correct lysine was ubiquitinated. We detect ubiquitin on FANCD2 K561 and FANCI K523 and do not detect any secondary ubiquitination events (Figure EV3). In addition, these reaction conditions also support full monoubiquitination of the isolated *Xenopus leavis* (*xl*) FANCD2 and FANCI substrates (Figure EV2). Purification profiles of both *hs*FANCD2-Ub and *hs*FANCI-Ub show no apparent changes in oligomeric state or hydrodynamic radius compared to non-ubiquitinated proteins (Figure 2C). Finally, we purified *hs*PCNA-Ub using a previously described method that uses a mutant E2 (UbcH5c S22R^25^) (Figure 2C). With the substrates purified (Figure 2C), we assessed whether *hs*FANCD2-Ub, *hs*FANCI-Ub or *hs*PCNA-Ub are directly deubiquitinated by USP1^FL^-UAF1 *in vitro*. The recombinant USP1^FL^-UAF1 is able to fully deubiquitinate *hs*FANCD2-Ub, *hs*FANCI-Ub and *hs*PCNA-Ub at sub-stoichiometric concentrations, in a UAF1 dependent manner (Figure 2D). In contrast, a catalytically-dead mutant of USP1 (C90S) is unable to deubiquitinate any of the substrates (Figure 2D). Taken together, these data establish that recombinant USP1-UAF1 is able to directly deubiquitinate diverse monoubiquitinated substrates *in vitro* without the need for auxiliary factors.

**Figure 2.**
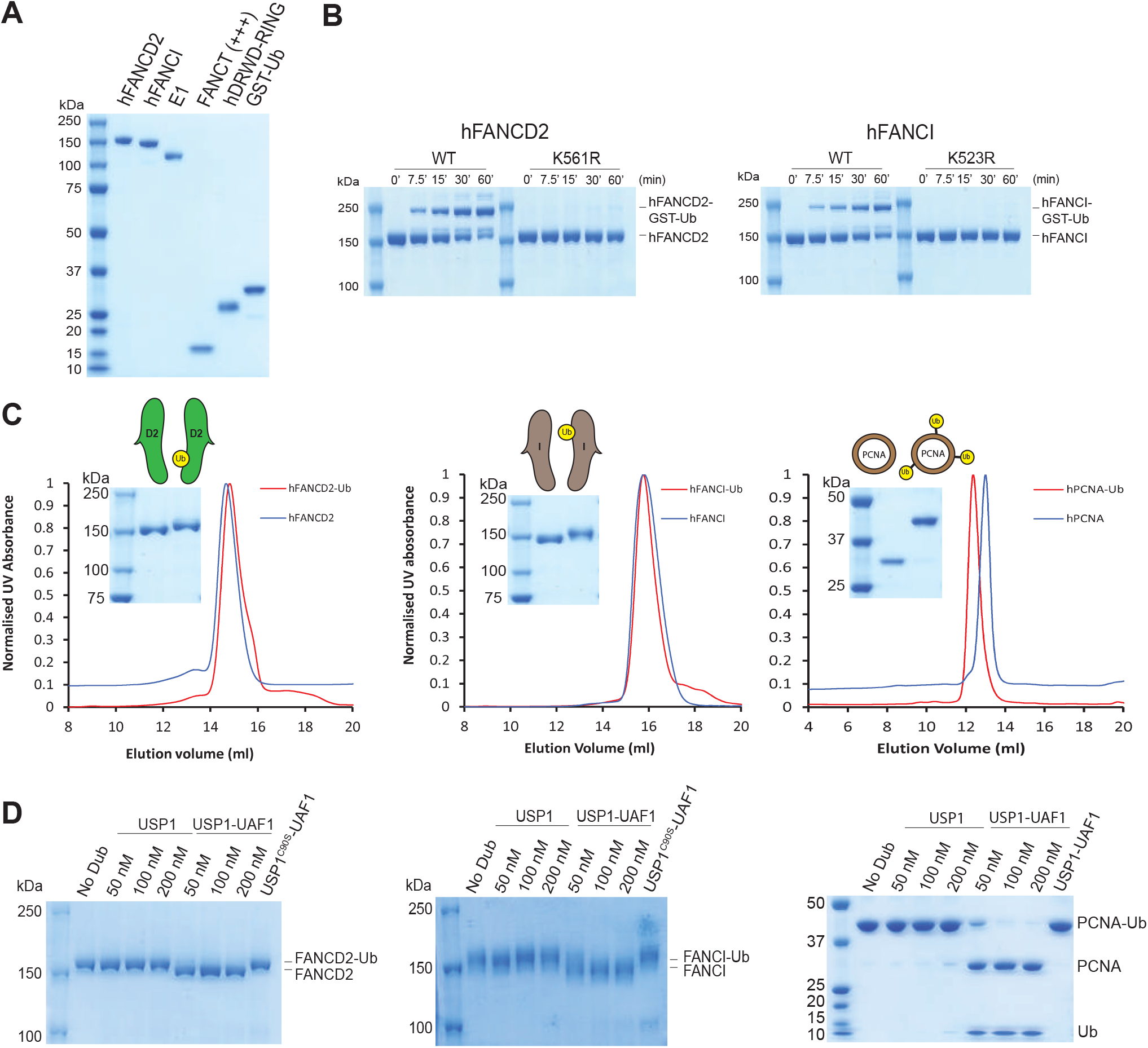
Recombinant *hs*FANCD2-Ub and *hs*FANCI-Ub is deubiquitinated by USP1-UAF1 *in vitro*. A) Purification of *hs*FANCD2 and *hs*FANCI, a hyperactive mutant Ube2T^V3^, FANCL^ΔELF^, ubiquitin E1 and GST-Ubiquitin. B) Ubiquitination reactions using reaction components in A. *hs*FANCD2 with K561R and FANCI with K523R mutations were used as a control for lysine specificity. C) Purity of monoubiquitinated *hs*FANCD2 and *hs*FANCI and size exclusion chromatography comparing modified and unmodified *hs*FANCD2 and *hs*FANCI. D) Deubiquitination of *hs*FANCD2-Ub, *hs*FANCI-Ub and *hs*PCNA-Ub by USP1 with and without UAF1.

**Figure EV2.**
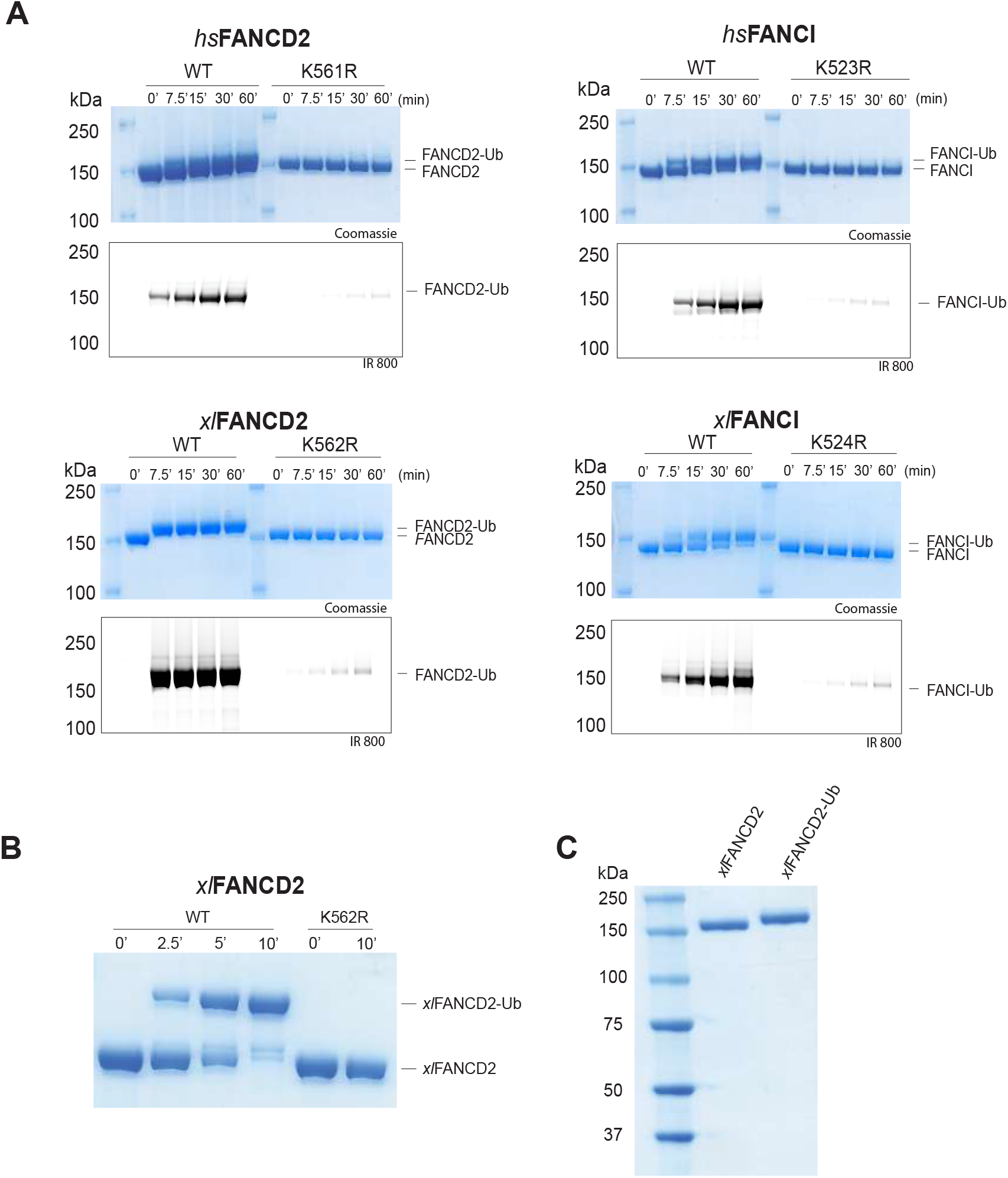
Purification of monoubiquitinated *Xenopus laevis* FANCD2. A) Monoubiquitination reactions of *hs*FANCD2, *hs*FANCI, *x*/FANCD2 and *x*/FANCI with fluorescent ubiquitin. Reactions used a hyperactive form Ube2T^V3^, ubiquitin, E1 and FANCL^ΔELF^. B) Ubiquitination of *x*/FANCD2 by GST-Ubiquitin with reaction conditions used above. C) SDS-PAGE, coomassie blue staining and western blots of recombinant *x*/FANCD2 and purfied monoubiquitinated *x*/FANCD2.

**Figure EV3.**
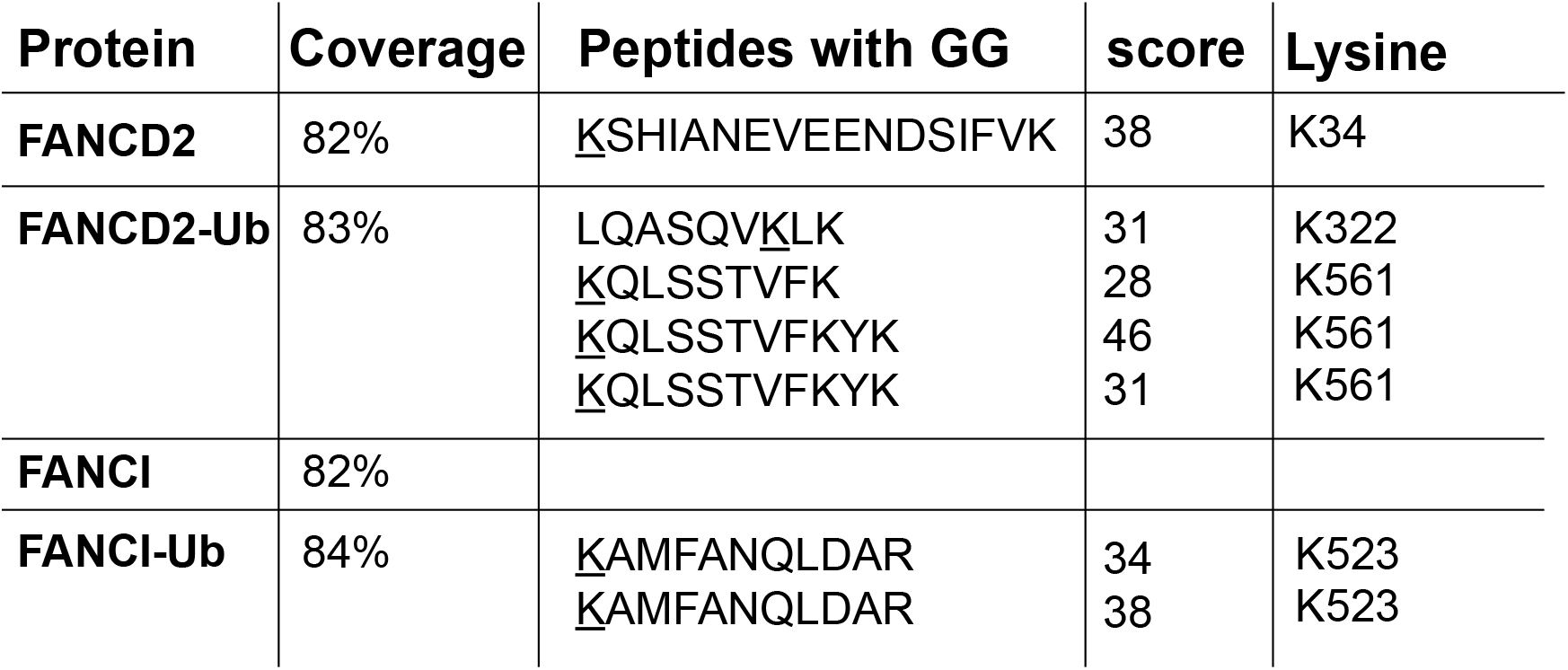
Ubiquitination sites detected by mass spectrometry on recombinant *hs*FANCD2, *hs*FANCD2-Ub, *hs*FANCI, and *hs*FANCI-Ub. Mass spectrometry on recombinant *hs*FANCD2 and *hs*FANCI proteins showing the amount of protein covered (%) and peptides which contain GG on a lysine residue, a Mascot score is shown for confidence.

### The N-terminus of USP1 is critical for FANCD2-Ub deubiquitination

To understand the minimum requirements for deubiquitination, we wondered whether the catalytic domain of USP1 is sufficient for deubiquitinating its physiological substrates. In order to test this, we assayed multiple USP1 fragments. As shown, USP1^FL^-UAF1 deubiquitinates *hs*FANCD2-Ub, *hs*FANCI-Ub and *hs*PCNA-Ub (Figure 3A). In addition, the catalytic module (USP1^ΔNΔ1Δ2^-UAF1) is sufficient at deubiquitinating *hs*FANCI-Ub (Figure 3A). Surprisingly, USP1^ΔNΔ1Δ2-^ UAF1 is unable to deubiquitinate *hs*FANCD2-Ub and there is slower activity on *hs*PCNA-Ub at early time points (Figure 3A), indicating the catalytic domain is not sufficient despite the catalytic domain being active on other substrates including *hs*FANCI. In light of this unexpected result, we wanted to determine which deleted region of USP1 is responsible for the loss in deubiquitinating activity on *hs*FANCD2-Ub. We therefore assayed each deletion: USP1^ΔN^, USP1^Δ1^ and USP1^Δ2^. Both deletions of insert 1 and 2 cleave at the same rate as USP1^FL^, however, deletion of the N-terminus results in a large loss of activity on *hs*FANCD2-Ub (Figure 3B). In contrast to *hs*FANCD2-Ub, none of the deletions have any effect on *hs*FANCI-Ub deubiquitination or *hs*PCNA-Ub deubiquitination (Figure 3B). Since the minimal USP1^ΔNΔ^1Δ2 is still an active deubiquitinase we wanted to determine whether at higher concentrations *hs*FANCD2-Ub could be deubiquitinated by this fragment. Full deubiquitination of *hs*FANCD2-Ub can be achieved by increasing the concentration of USP1^ΔNΔ1Δ2^ to a 1:1 ratio (Figure 3C). This is in contrast to USP1^Δ1Δ2^, which is able to deubiquitinate *hs*FANCD2-Ub at sub-stoichiometric concentrations (Figure 3C). These data indicate that although the N-terminus is not required for catalytic activity *per se*, the N-terminus contributes to a much more productive *hs*FANCD2-Ub deubiquitinase.

**Figure 3.**
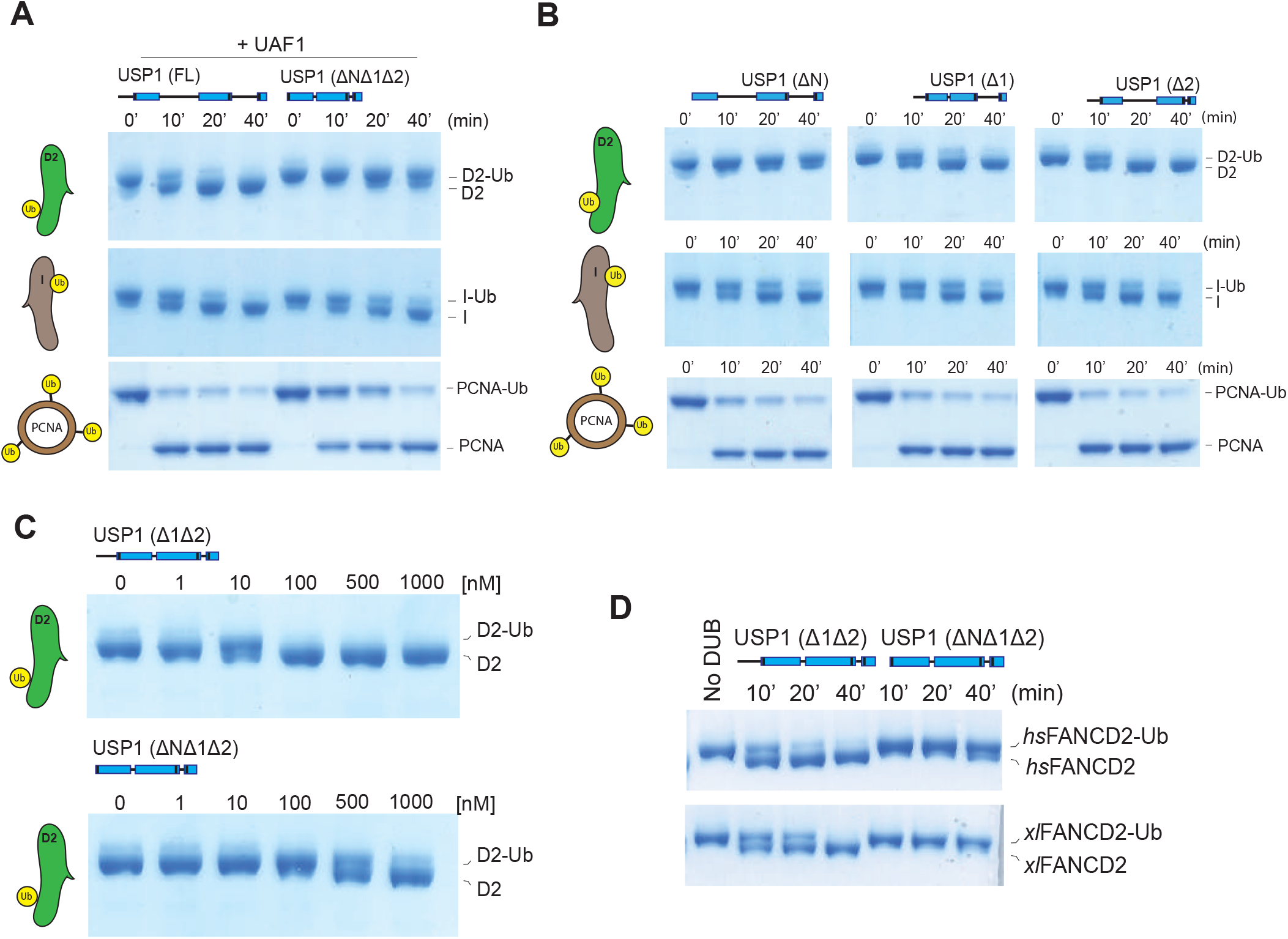
The N-terminus of USP1 is required for robust FANCD2-Ub deubiquitination but not for *hs*FANCI-Ub or *hs*PCNA-Ub. A) Deubiquitination reactions of recombinant *hs*FANCD2-Ub, *hs*FANCI-Ub sand *hs*PCNA-Ub using USP1^FL^-UAF1 and USP1^ΔNΔ1Δ2^-UAF1. B) Deubiquitination assays of *hs*FANCD2-Ub, *hs*FANCI-Ub and *hs*PCNA-Ub using indicated USP1 truncations. C) Full deubiquitination of *hs*FANCD2 requires 10x USP1^ΔNΔ1Δ2^ when compared to USP1^Δ1Δ2^. D) Deubiquitination of frog (*Xenopus leavis*) FANCD2 by USP1^Δ1Δ2^ and USP1^ΔNΔ1Δ2^. Proteins are separated by SDS-PAGE and stained with coomassie blue.

A recent report suggests that *x*/FANCI is required for human USP1-UAF1 to remove ubiquitin from *x*/FANCD2-Ub^32^. In contrast, we find that *hs*FANCD2 does not require the presence of *hs*FANCI in order to be deubiquitinated. Therefore, we wondered if our observation was due to the use of *hs*FANCD2 substrate. In order to test this, we purified *x*/FANCD2-Ub to homogeneity (Figure EV2) and assayed with human USP1-UAF1. In contrast to previous reports, we find that USP1^FL^ and USP1^Δ1Δ2^ are able to fully deubiquitinate *x*/FANCD2-Ub (Figure 3D). Furthermore, consistent with our observations using *hs*FANCD2-Ub as substrate, deletion of the N-terminus reduces USP1 activity for *x*/FANCD2-Ub (Figure 3D). Since deletion of the N-terminus of USP1 does not result in a defect for Ub-KG^TAMRA^ (Figure 1E), K63/K48 diUb (Figure 1F), *hs*FANCI-Ub (Figure 3B) and *hs*PCNA-Ub (Figure 3B), it is likely that the N-terminus is a specific FANCD2-Ub requirement that is extended from the catalytic domain. Taken together these data suggest that the N-terminus of USP1 harbours a substrate targeting sequence specific for FANCD2.

### The N-terminus drives specific FANCD2 K561-Ub deubiquitination

The ubiquitination site of FANCD2 is at a specific residue, K561. We wondered whether the FANCD2 targeting by the N-terminus of USP1 also extends to the site of ubiquitination i.e. K561-Ub. To test this we generated FANCD2 and modified lysines distinct from K561 (Figure 4A). We decided to use a mutant E2 (UbcH5c^S22R^) which primarily monoubiquitinates proteins^25^. As monoubiquitination activity was weak, we used the E3 ligase RNF4 fragment (RNF4-RING fusion, RNF4^RR^) to increase the activity^34^. The UbcH5c^S22R^ and RNF4^RR^ pair resulted in robust and multiple-ubiquitination events on *hs*FANCD2 (Figure 4B). To ensure K561 is not being monoubiquitinated by UbcH5c^S22R^and RNF4^RR^, we used K561R *hs*FANCD2 as the substrate. We performed an E3 assay to monoubiquitinate *hs*FANCD2 and *hs*FANCD2 K561R, which we term *hs*FANCD2 K561-Ub and *hs*FANCD2 KX-Ub, respectively (Figure 4A), and treated the reaction with USP1^FL^-UAF1 or USP1^ΔN^UAF1. Interestingly, while USP1^FL^-UAF1 deubiquitinates both substrates K561-Ub and KX-Ub at similar rates, the USP1^ΔN^ –UAF1 shows clear activity on KX-Ub but little detectable activity on K561-Ub (Figure 4C and 4D). These data show that the USP1 N-terminus targeting sequence also specifies the site for deubiquitination of FANCD2-Ub i.e. K561.

**Figure 4.**
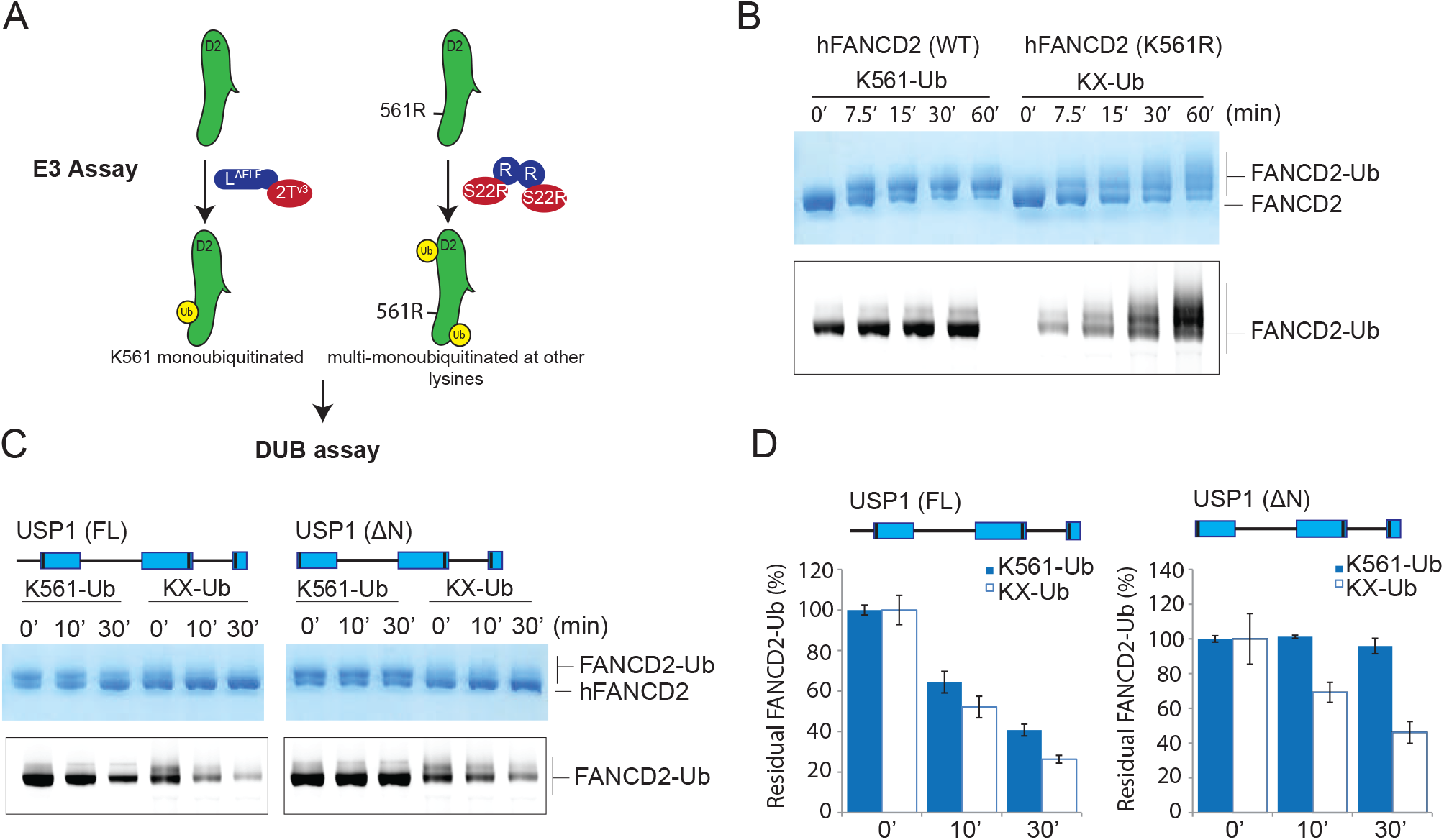
The N-terminus of USP1 drives specifc K561 deubiquitination of FANCD2. A) Schematic of *hs*FANCD2 monoubiquitination reactions to make monoubiquitinated FANCD2-K561-Ub and FANCD2-KX-Ub (where X denotes random lysines). B) Monoubiquitination of *hs*FANCD2 (WT) by Ube2T^V3^ and FANCL^ΔELF^ is predominantly a single monoubiquitination event at K561. Monoubiquitination of *hs*FANCD2 K561R by UbcH5c (S22R) and RNF4 (RNF4-RING fusion) results in multi-monoubiquitination of other lysines of FANCD2. Ubiquitinated products are formed within a time dependent manner as shown by coomassie blue staining and fluroescently labelled ubiquitin, which only shows ubiquitinated products. C) USP1^FL^-UAF1 deubiquitinated FANCD2 regardless of the ubiquitin’s location. USP1^ΔN^-UAF1 deubibiquitinates other lysines much faster than K561-Ub. Monoubiquitinated FANCD2-K561-Ub or FANCD2-KX-Ub reactions were arrested by apyrase and subject to USP1^FL^-UAF1 or USP1^ΔN^-UAF1. Fluorescent ubiquitin is used to monitor ubiquitinated species and quantify DUB activity. Data shows intensities as percent of ubiquitinated input [n = 3] +/− SD.

### The N-terminus of USP1 contains critical residues for FANCD2-Ub deubiquitination

In order to identify residues within the N-terminus of USP1 required for *hs*FANCD2-Ub specificity, we looked at regions that are well conserved (Figure 5A). The N-terminus of USP1 is almost completely conserved from humans to zebrafish between residues 19–40 (Figure 5A). To screen mutants of USP1, we optimised and purified USP1^Δ^1Δ2 and N-terminus variants from *E.coli* (Figure EV4). Similar to deleting residues 1–54 of USP1, deletion of residues 21–29 also resulted in a specific loss in activity for *hs*FANCD2-Ub but not *hs*PCNA-Ub (Figure 5B). We next determined whether any of the side chains are important, so we mutated each residue to alanine and assayed for DUB activity using Ub-prg, *hs*PCNA-Ub, *hs*FANCI-Ub and *hs*FANCD2-Ub (Figure EV4 and EV5). All point mutants are active with Ub-prg, *hs*PCNA-Ub and *hs*FANCI-Ub, in contrast, there is a loss in activity for several point mutants N21A, R22A, L23A, S24A and K26A on *hs*FANCD2-Ub deubiquitination (Figure EV5). To quantify the relative differences between USP1 mutants we purified *hs*FANCD2-Ub with a fluorescently labelled Ub^800^ that allows us to detect only the ubiquitinated FANCD2 (*hs*FANCD2-Ub^800^) (Figure EV6). Using this approach we monitored the deubiquitination of *hs*FANCD2-Ub^800^ by each mutant and quantified the remaining FANCD2-Ub as a percentage of the input (Figure 5C). We find that while USP1^Δ^1Δ2-UAF1 cleaves ∼80% of the *hs*FANCD2-Ub^800^, USP1 ^ΔNΔ1Δ2^-UAF1 shows no apparent cleavage (Figure 5C). Interestingly, R22A and L23A were able to only deubiquitinate ∼10% of the *hs*FANCD2-Ub^800^ (Figure 5C). We did not further characterise L23A as it co-purifies with a large contaminant chaperone, which may suggest some misfolding (Figure EV4). We next wondered whether the charge at R22 is critical for deubiquitination as it is conserved as a lysine in zebrafish (Figure 5A). We generated R22K, R22A and R22E variants. As expected, R22A, R22E and R22K have no effect on *hs*PCNA-Ub or *hs*FANCI-Ub deubiquitination (Figure 5D). In contrast, we observe that R22A has an intermediary affect and R22E is comparatively more compromised on *hs*FANCD2-Ub deubiquitination (Figure 5D). R22K retains WT activity on hFANCD2-Ub (Figure 5D), suggesting a positive charge is required. Finally, we showed in the full length USP1-UAF1 context, a single point mutation in its N-terminus (R22E) is sufficient for specifically disrupting *hs*FANCD2-Ub deubiquitination but not *hs*FANCI-Ub or *hs*PCNA-Ub (Figure EV7). Taken together, these data indicate that R22 plays an important role in deubiquitination of *hs*FANCD2-Ub. However, it is clear that multiple conserved residues (N21, R22, L23, S24 and K26), that are within a short N-terminal region, drive USP1 mediated FANCD2-Ub deubiquitination.

**Figure 5.**
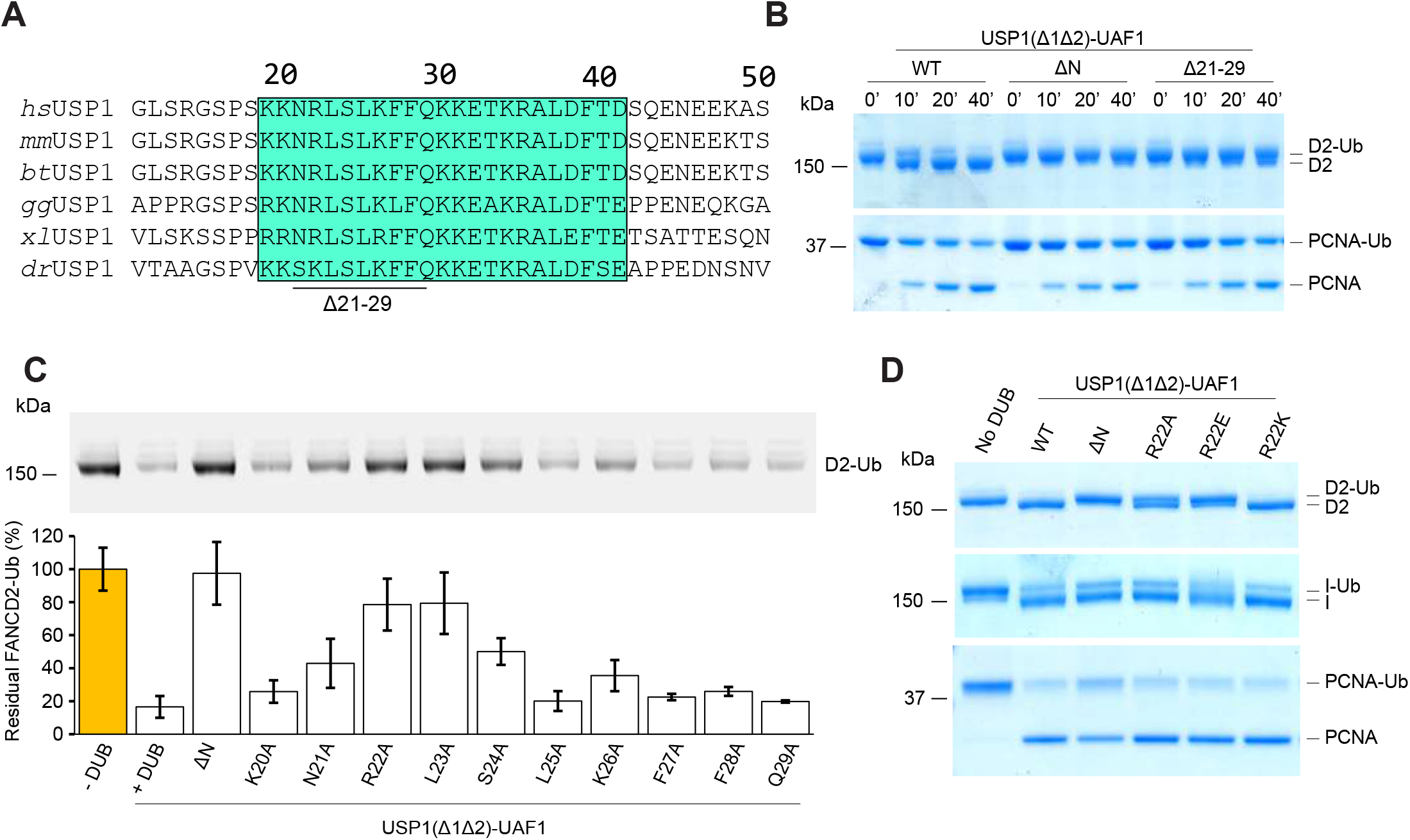
Several highly conserved residues within USP1 N-terminus determine FANCD2-Ub deubiquitination. A) Sequence alignment of USP1 N-terminus from several species (*hs: Homo sapiens, mm: Mus musculus, bt: Bos taurus, gg: Gallus gallus, xl: Xenopus leavis, dr: Danio rerio*). Deleted USP1 residues in B) are indicated. Deubiquitination assay of *hs*FANCD2-Ub and *hs*PCNA-Ub using *E.coli* purifed USP1^Δ1Δ2^, USP1^ΔNΔ1Δ2^ and USP1^Δ21^-29Δ1Δ2 in complex with UAF1 C) Deubiquitination assays of hFANCD2-Ub^800^ by recombinant USP1^Δ1Δ2^ Ala point mutants from *E.coli*. Alanine point mutations within the USP1 N-terminus result in a loss of FANCD2-Ub^800^ deubiquitination. The loss in fluorescence intensity was quanitified and normalised to the amount of input hFANCD2-Ub^800^. Data displays mean residual FANCD2-Ub^800^ (%) [n = 3] +/− SD normalised to input. D) R22 point mutations indicate that a positive charge at position 22 is critical for *hs*FANCD2-Ub deubiquitination but not for *hs*FANCI-Ub or *hs*PCNA-Ub.

**Figure EV4.**
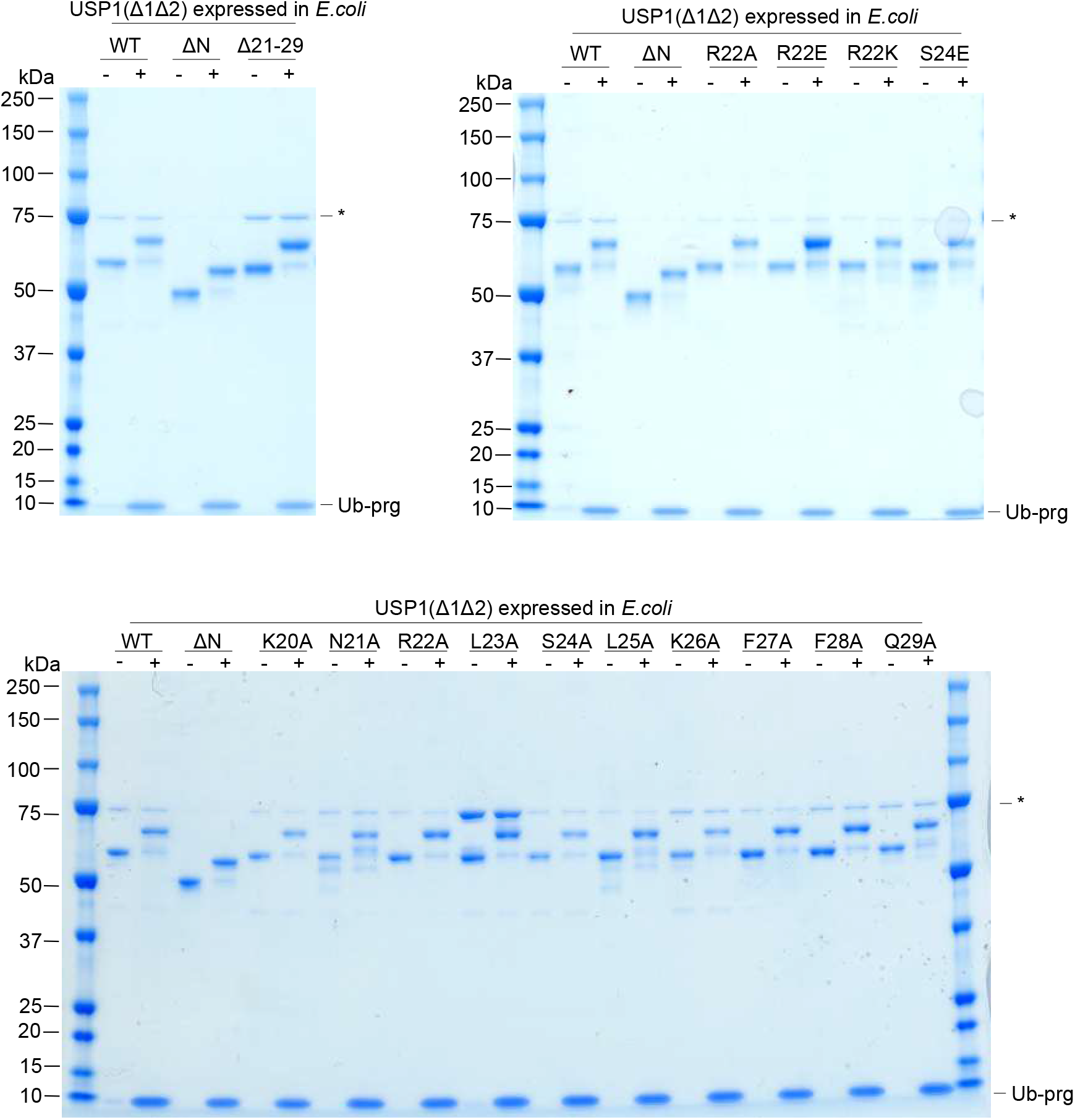
Recombinant USP1 catalytic domain with and without N-terminus expressed in *E.coli* and reacted with Ub-prg. SDS-PAGE and coomassie staining of USP1^Δ1Δ2^ and point mutants within the N-terminus. Excess Ub-prg reacts with all mutants of USP1, as shown by a slower migrating band, indicating proper folding of the catalytic domain.

**Figure EV5.**
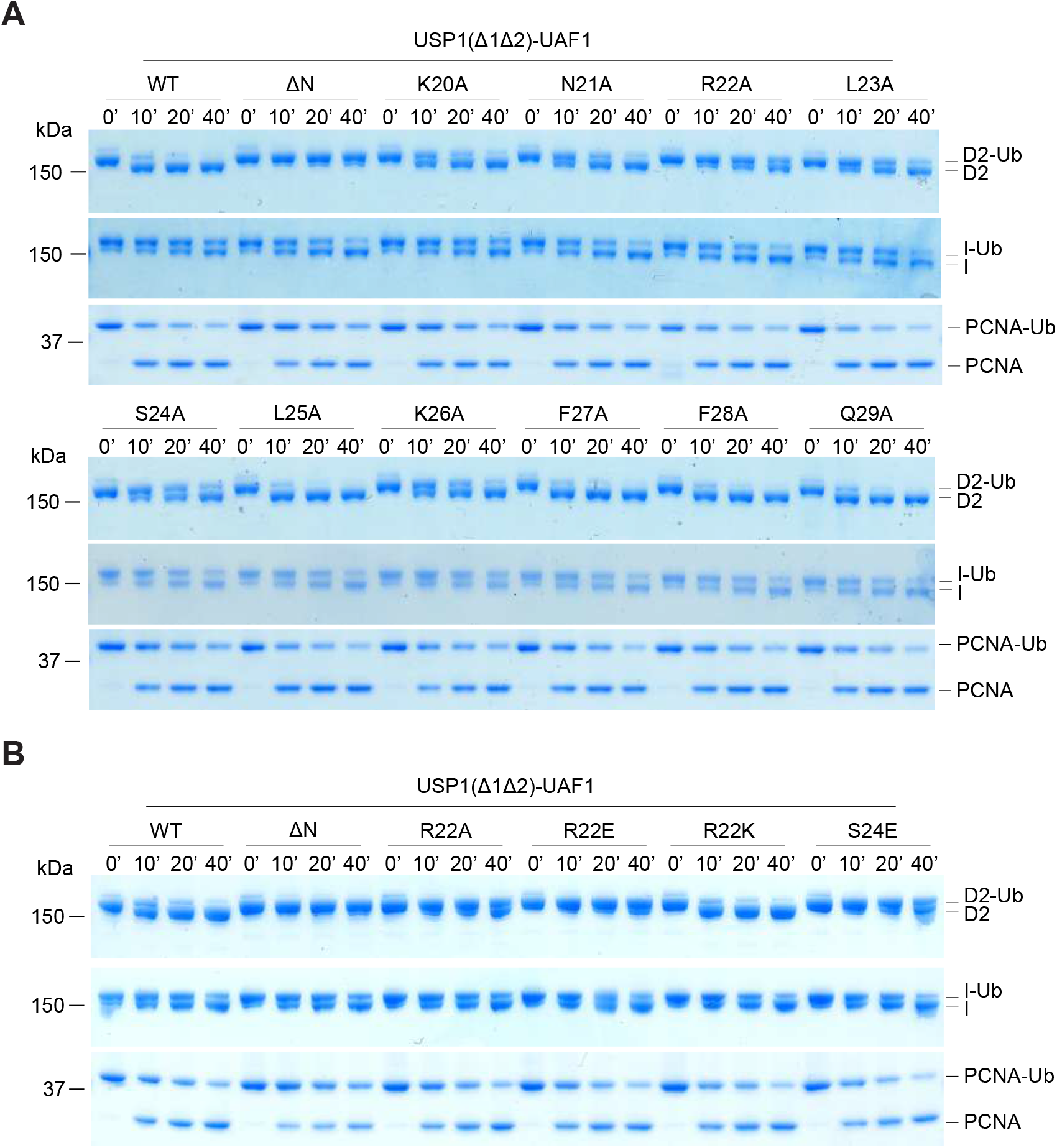
Deubiquitination of hsFANCD2-Ub, hsFANCI-Ub and hsPCNA-Ub by *E.coli* expressed USP1^Δ1Δ2^ and point mutants. A) Deubiquitination reactions of recombinant *hs*FANCD2-Ub, *hs*FANCI-Ub sand *hs*PCNA-Ub using USP1^Δ1Δ2^ and N-terminus point mutants expressed in *E.coli*. USP1 is complexed with equimolar UAF1. B) Deubiquitination assays of substrates by USP1 with indicated mutants.

**Figure EV6.**
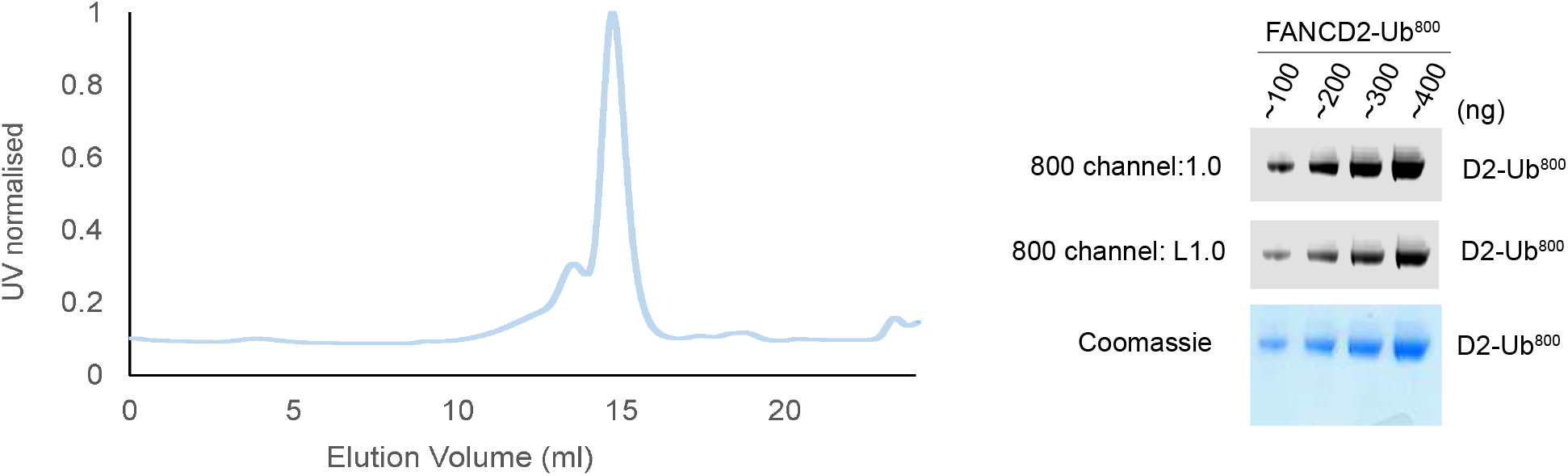
Purity of FANCD2 monoubiquitinated with ubiquitin^800^. Size exclusion chromatography of *hs*FANCD2-Ub^800^ and SDS-PAGE analysis stained with coomassie blue and scanned with licor 800 channel, which shows only ubiquitinated FANCD2.

**Figure EV7.**
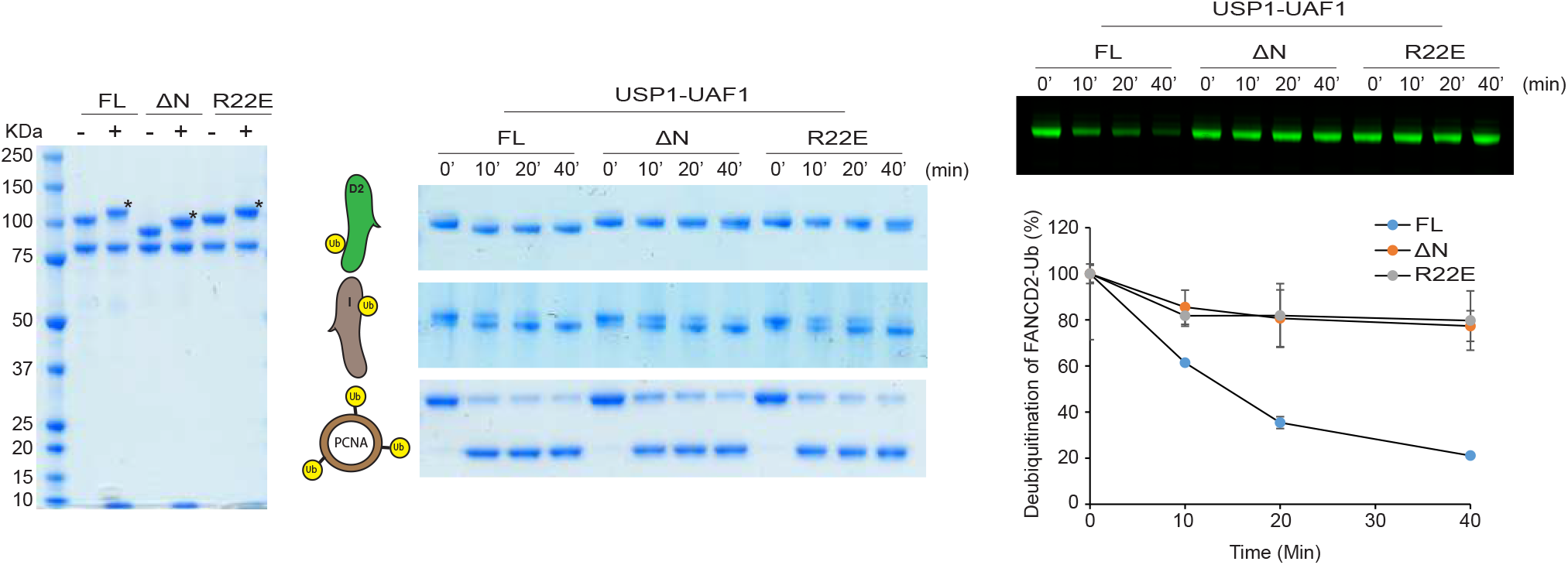
A single point mutation within the N-terminus of USP1 can disrupt deubiquitination of hsFANCD2 but not hsFANCI or hsPCNA. A) SDS-PAGE analysis of recombinant USP1, USP1^ΔN^ and USP1^R22E^ in complex with UAF1 with and without Ub-prg. B) Deubiquitination of recombinant *hs*FANCD2-Ub, *hs*FANCI-Ub and *hs*PCNA-Ub by proteins in part A. C) Deubiquitination of FANCD2-Ub^800^ by proteins in A. Data shows deubiquitination as mean percentage over time [n = 2] +/− SD. 100% is set as 0’ time point.

### The N-terminus of USP1 is a FANCD2 recognition element

Mutation or deletion of the USP1 N-terminus specifically and negatively impacts FANCD2-Ub deubiquitination. We hypothesised that this region in the N-terminus of USP1 encodes a FANCD2 binding site. To test this, we purified UAF1 with an N-terminal Strep tag and formed stable complexes with USP1^Δ1Δ2^ or USP1^ΔNΔ1Δ2^ harbouring C90S mutations in order to disrupt catalytic activity. We assayed interaction with *hs*FANCD2-Ub and find that USP1^Δ1Δ2^-UAF1 complexes can capture *hs*FANCD2-Ub (Figure 6A). In contrast, assaying capture with USP1^ΔNΔ1Δ2^-UAF1, we detect little *hs*FANCD2-Ub capture on beads, when compared to USP1^Δ1Δ2^ (Figure 6A). Since the interaction is defective but not absent with USP1^ΔNΔ1Δ2^ when using *hs*FANCD2-Ub, we wondered whether the ubiquitin on *hs*FANCD2 is required for the N-terminus to interact. We find that unmodified *hs*FANCD2 interacts with USP1^Δ^1Δ2 (Figure 6A). In contrast, we do not detect capture of *hs*FANCD2 when using USP1^Δ^NΔ1Δ2 as bait (Figure 6A). These data show that USP1 interacts with *hs*FANCD2 regardless of ubiquitination, but shows that the interaction is fully dependent on the N-terminus. Since we detect an interaction with USP1^ΔNΔ1Δ2^ with *hs*FANCD2-Ub, albeit much less than with the N-terminus, this suggests a ubiquitin-mediated interaction, likely through the catalytic domain of the USP1 (Figure 6A).

As USP1’s interaction with *hs*FANCD2 is dependent on the N-terminus, we wanted to determine whether this is specific for *hs*FANCD2, so we similarly assayed interactions of *hs*FANCI and *hs*PCNA. Interestingly, we can detect an interaction with *hs*FANCI and *hs*FANCI-Ub, but that is unaffected by deletion of the N-terminus of USP1 (Figure EV8). We could not detect an interaction between USP1 and *hs*PCNA, but we observe interaction with *hs*PCNA-Ub (Figure EV8). Which again likewise and consistent with deubiquitination assays, is not dependent on the N-terminus of USP1. Together this data indicates that the N-terminus of USP1 specifically interacts with *hs*FANCD2 regardless of ubiquitination. We do note an enhancement in interaction with *hs*FANCD2-Ub when compared to *hs*FANCD2 and thus suggest that there are two points of interaction; FANCD2: N-term and USP domain: ubiquitin.

**Figure 6.**
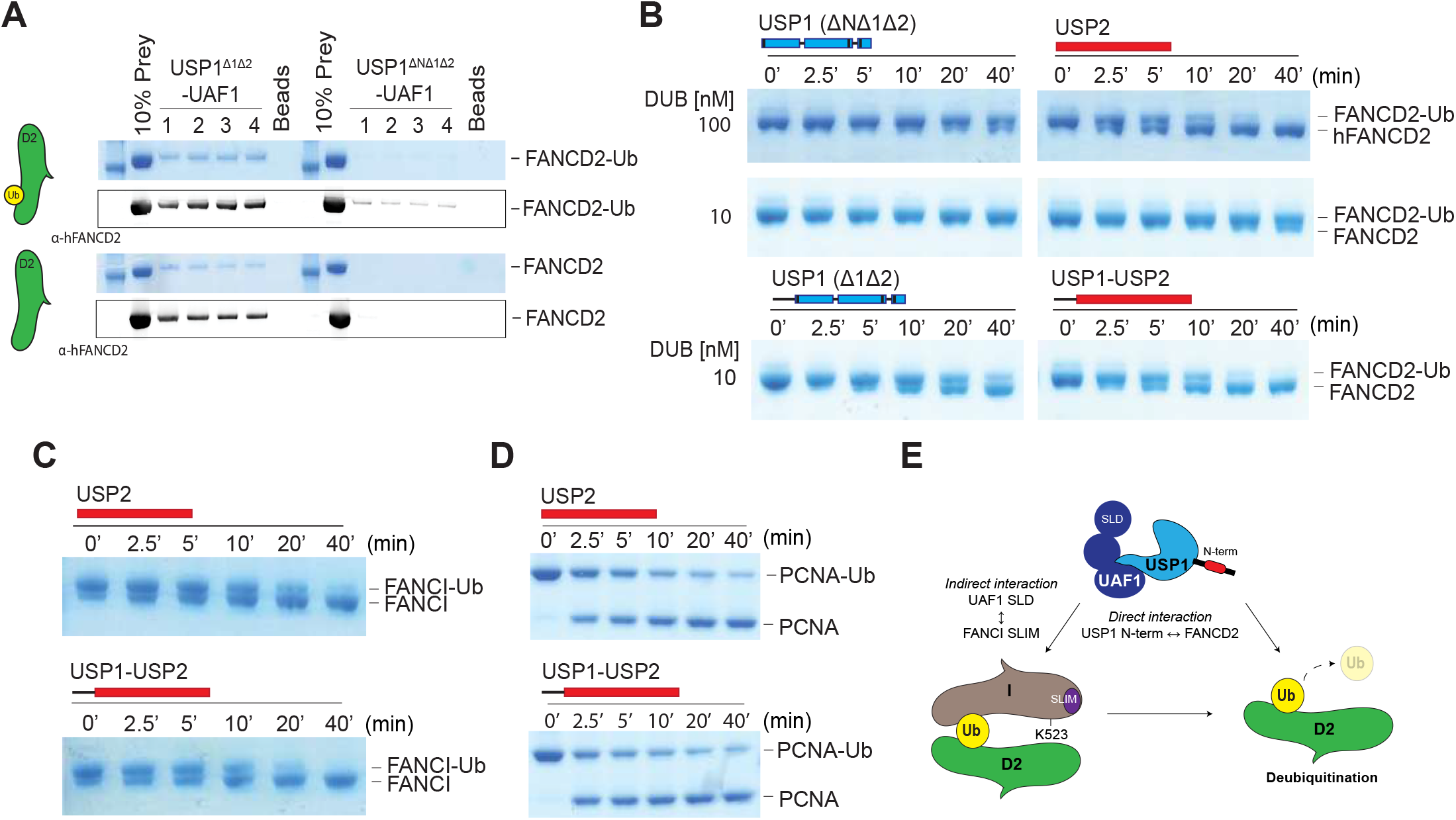
The N-terminus of USP1 binds to FANCD2 and can enchance USP2 deubiquitinase activity against FANCD2 when fused N-terminal to the catalytic domain. A) Pull down capture of *hs*FANCD2-Ub and *hs*FANCD2 by strep-tagged UAF1 bound to USP1^Δ1Δ2^ USP1^ΔNΔ1Δ2^. Data shows western blots with [n = 4] technical replicates, experiment was performed twice. B) Deubiquitination of *hs*FANCD2-Ub by USP2 cataytic domain and an N-terminal fusion of USP1 N-terminus to USP2 catalytic domain (USP1^1^-60-USP2) at two different concentrations. C) *hs*FANCI-Ub and D) *hs*PCNA-Ub deubiquitination by USP2 and USP1^1^-60-USP2 shows no apparent difference in activity indicating that USP1 N-terminus fused to the catalytic domain of USP2 provides a FANCD2-Ub specific gain-in-activity. E) Model of FANCD2-Ub deubiquitination by USP1-UAF1. USP1 targets FANCD2 via a direct interaction using its N-terminus. In addition to the N-terminus of USP1, FANCD2-Ub/FANCI complexes can be indirectly targeted by an interaction between UAF1 Sumo-like domain (SLD) and FANCI sumo-like interacting motif.

**Figure EV8.**
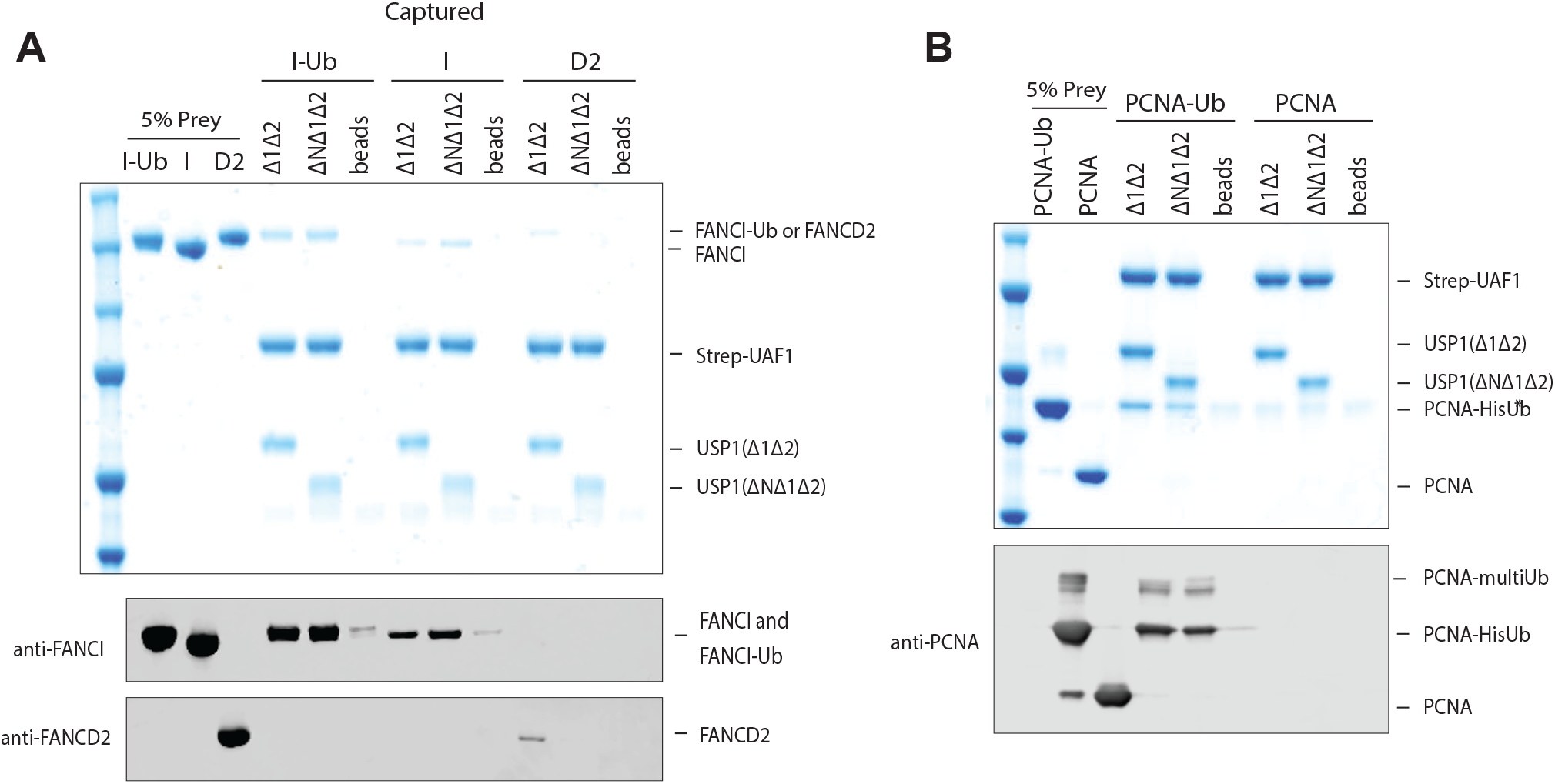
Interaction of USP1 with *hs*FANCI and *hs*PCNA do not depend on N-terminus of USP1. A) Strep-tag immobilised UAF1 bound to USP1^Δ1Δ2^ captures *hs*FANCI and *hs*FANCI-Ub, however, deletion of the N-terminus (USP1^ΔNΔ1Δ2^) did not abolish capture. B) Immobilised UAF1 bound to USP1^Δ1Δ2^ or USP1^ΔNΔ1Δ2^captures *hs*PCNA-Ub. However, *hs*PCNA could not be captured under these conditions.

### A chimera fusion of USP1 N-terminus to USP2 catalytic domain provides FANCD2 specificity

Deletion of the N-terminus of USP1 does not appear to affect the catalytic activity of the catalytic module (USP1^ΔNΔ1Δ2^), except in the case of *hs*FANCD2-Ub deubiquitination. We therefore wondered whether the N-terminus can increase the efficiency of FANCD2-Ub deubiquitination of a distinct DUB module. To test this, we chose the catalytic domain of USP2, generally considered promiscuous deubiquitinase due to its lack of substrate specificity^35^. We created and purified a chimeric fusion USP1^1^-60-USP2 where residues 1–60 of USP1 are N-terminal to the USP2 catalytic domain (Figure 6B). We find that USP2 at 100 nM is able to completely deubiquitinate 1µM *hs*FANCD2-Ub by 40 min (Figure 6B), in contrast, the chimeric USP1^1^-60-USP2 achieves this in 10 min (Figure 6B). Since the N-terminus increases USP2 activity against *hs*FANCD2-Ub, we asked whether the N-terminus increases activity of USP2 towards other ubiquitinated substrates. In contrast to *hs*FANCD2-Ub, there are no differences in activity of USP2 and the USP1^1^-60-USP2 chimera on either *hs*FANCI-Ub (Figure 6C) or *hs*PCNA-Ub (Figure 6D). These data show that the addition of the N-terminus in *cis* confers a gain-of-activity to USP2. The N-terminus of USP1 contains a FANCD2 determinant sequence that functions at the core of FANCD2 deubiquitination by specifically binding and increasing deubiquitination efficiency (Figure 6E).

## Discussion

A cycle of monoubiquitination and deubiquitination of FANCD2 is critical for completion of FA ICL repair^8^. Removal of the ubiquitin signal from FANCD2 is conducted by the USP1-UAF1 deubiquitinase, and disruption of USP1 catalytic activity results in an accumulation of FANCD2-Ub and FA-like phenotypes^6,9,11,23^. Despite multiple DUBs existing at DNA replication and repair foci, USP1 appears to be the only DUB whose loss results in an accumulation of FANCD2-Ub^6^. The molecular determinants and mechanisms of how DUBs such as USP1 target and regulate distinct pools of substrates have remained elusive, due to the lack of molecular insights into the process. Here we report a unique feature at the N-terminus of USP1 and a potential short linear interacting motif (SLIM) at the N- terminus explains that it specifically and directly targets K561 of FANCD2, providing a molecular foundation for how USP1 targets one of its substrates with high specificity.

USP1 activity is subject to layers of regulation during cell-cycle progression. For example, USP1 is turned over throughout the cell cycle, activated and stabilised by binding partners such as UAF1 and, in addition, undergoes autocleavage to allow targeting by both the N-end rule and C-end rule pathways^7,11,22,36^. However, we show that USP1, in complex with UAF1, deubiquitinates at least three DNA repair related substrates *in vitro*; FANCD2-Ub, FANCI-Ub and PCNA-Ub. Thus, this suggests everything essential for deubiquitination is encoded within USP1-UAF1. Interestingly, UAF1 is an abundant protein with multiple functions including activating USP12 and USP46^19^. It has been suggested that UAF1 may indirectly target USP1 to specific substrates such as FANCD2 or PCNA via a C-terminal sumo-like domain (SLD) that binds to sumo-like interacting motifs on protein partners of FANCD2 and PCNA^24^ (Figure 6E). However, disruption of either USP12 or USP46 does not affect FANCD2-Ub levels, showing that FANCD2 is not an overlapping substrate^19^. Since the SLD would target multiple DUBs to the same substrates, further layers of regulation may exist to ensure the correct DUB is targeted to FANCD2-Ub or PCNA-Ub. By reconstituting FANCD2-Ub deubiquitination by USP1-UAF1, we identified the minimal components required for deubiquitination and discover a unique sequence within the N-terminus of USP1 critical for removing ubiquitin from FANCD2 (Figure 6E). The N-terminus bound specifically to FANCD2, even in the absence of ubiquitin, and did not bind to PCNA or FANCI. Thus USP1 displays a higher level of substrate targeting for FANCD2. It is interesting that the N-terminus of USP1 is not required for FANCI-Ub deubiquitination, since FANCI shares a similar ubiquitination site to FANCD2 both structurally and on a sequence level, allowing us to speculate that perhaps FANCD2 is the major substrate of USP1. USP1 targets multiple substrates, so it will be exciting to determine what further features USP1 or even the substrates have that may contribute to substrate targeting and catalysis, either through the N-terminus or insertions of USP1 or even unique features of the substrates.

It is interesting that activity of USP1 is lost for FANCD2-Ub deubiquitination when deleting the N-terminus but not for the other substrates. This specific loss in activity suggests that USP1 without the N-terminus may struggle to form a productive enzyme-substrate complex with FANCD2. The N-terminus of USP1 is an extension of the USP domain and is ∼75 amino acids in length. It is unclear how flexible, structured or compact the N-terminus may be in relative position to the USP domain and catalytic site. Therefore, it is difficult to predict whether the N-terminus of USP1 directly contributes to catalysis when FANCD2 is bound, or simply acts to recruit FANCD2-Ub and stabilise the isopeptide bond within the active site. Since USP1^ΔN^UAF1 loses activity for ubiquitinated K561, but not for other lysines of FANCD2, this allows us to speculate that the N-terminus binds and precisely positions FANCD2 in order to increase activity for a specific lysine position. It can also be suggested that FANCD2-K561-Ub is not in a fully accessible conformation for deubiquitination, and that the N-terminus modulates this conformation to allow formation of a more productive complex. This is reminiscent of a recently identified monoubiquitination site of SETDB1, which protects the ubiquitin from deubiquitination via multiple SETDB1-Ubiquitin interactions^37^, perhaps FANCD2 also partially protects and occludes ubiquitin at K561, and the N-terminus is required to relieve this effect. It will be interesting to determine how the N-terminus binds to FANCD2, how it modulates USP1 function towards FANCD2 and whether other PTMs or processes within the N-terminus can modulate its function.

It is not understood how the majority of USPs maintain specificity as most of them target a common factor, ubiquitin. Interestingly, USPs have distinct sets of substrates as individual knockdown of DUBs will only affect certain substrates^6^. For some USPs they target a specific signal, for example, USP18 has a unique hydrophobic patch within the USP domain which targets a unique hydrophobic patch to the ubiquitin-like protein ISGN15^38^. Other examples include USP30, which shows preference for K6 linked ubiquitin chains via several residues in the USP domain^17,39^. However, here we have an example, USP1, which we have shown targets a specific ubiquitinated substrate, and therefore recognise both ubiquitin and the substrate itself. A similar example to this could be monoubiquitinated lysines on histones, which are commonly targeted for DNA repair signalling events. For example, Histone 2B is monoubiquitinated on K120 and targeted by the SAGA DUB module, a four protein complex which contains USP22 (Ubp8 homolog in yeast). Interestingly, while Ubp8 showed contributions for targeting nucleosomes via several residues, other members of the complex such as sgf11 made more significant contributions via the acidic patch on H2A/H2B, and therefore likely governs most of the substrate specificity^27^. While SAGA-DUB achieves specificity via a protein complex, other DUBs such as USP7 contain substrate targeting regions within the same polypeptide. USP7 displays substrate targeting regions via its N-terminal TRAF domain for p53^40^.

However, in contrast to the effect we see with disruption of USP1’s N-terminus on FANCD2-Ub deubiquitination, the N-terminus of USP7 only had a minor effect on p53 deubiquitination *in vitro*, indicating it is not critical for forming a productive enzyme-substrate complex^41^. We show that USP1 specifically targets one ubiquitinated lysine made by a specific E3 ligase, FANCL. Another DUB, USP48, was recently published to also specifically oppose the monoubiquitination made by another lysine specific E3 ligase BRCA1^26^. This suggests that some DUBs and E3 activities may have co-evolved to oppose each other on specific cellular signals and maintain an appropriate equilibrium of non-ubiquitinated and ubiquitinated substrate.

Arguably the most important known function of USP1 is to regulate FANCD2, as the monoubiquitinated form is important for both ICL repair and also origin of replication firing in latent conditions^42^. In addition, non-ubiquitinated FANCD2 is important in DNA surveillance mechanisms^43^, meaning the presence of both FANCD2-Ub and FANCD2 is important for DNA integrity and cell proliferation. Moreover, USP1 depletion increases oncogene-induced senescence (OIS) and plays a pivotal role in protecting genomic instability by preventing FANCD2-Ub aberrant aggregation^12^. USP1-UAF1 therefore works to recycle and maintain appropriate levels of both monoubiquitinated and un-ubiqutinated FANCD2. Inhibition of USP1-UAF1 activity by the specific small molecule, ML323, leads to an increased sensitivity to platinum based compounds in resistant cells^11^. Furthermore, the USP1-UAF1 complex has been shown to play critical roles in homologous recombination^10^. This collection of studies reveals that USP1 would be an excellent anti-cancer target by disrupting its ability to deubiquitinate FANCD2-Ub, as this would lead to OIS and cellular sensitivity to crosslinking agents^8,9^. However, USP1-UAF1 targets multiple substrates including ID2 proteins and TBK1, meaning an all-around catalytic activity inhibitor may affect multiple processes, perhaps a more specific inhibitor that could block FANCD2-USP1 interaction, such as the N-terminus, would provide a more specific therapy.

## Materials and Methods

### Cloning

Full length USP1^FL^ with G670A/G671A (mutated autocleavage site^7^) was ligated into a pFASTBAC vector with a TEV cleavable His tag by BamH-Not1 restriction cloning. Subsequent, truncations of USP1 were made in the full length background by PCR mutagenesis: N-terminus (Δ1–54), Insertion 1 (Δ229–408) and Insertion 2 (Δ608–737). UAF1 was ligated into a pFASTBAC vector with an N-terminal His-3C tag using BamH1-Not1, mutagenesis was used to add a Strep11 tag and a linker ‘GGSGGS’ N-terminal of the His-tag. *Xenopus leavis* FANCD2 and *Xenopus leavis* FANCI genes were a kind gift from Puck Knipscheer (Hubrecht Institute, Utrecht), genes were ligated into pFASTBAC vectors using EcoRI-XhoI and mutated to contain N-terminal His-TEV-HA and His-TEV-V5 tag, respectively. Human FANCD2 gene was synthesised by GeneArt Gene Synthesis (Invitrogen), expression optimised for *Spodoptera frugiperda* and subsequently cloned into a pFASTBAC vector with a His-3C tag using BamH1-Not1.

USP1^Δ1Δ2^ expression optimised for *E.coli* expression was synthesised by GeneArt Gene Synthesis (Invitrogen) and subsequently ligated into a pGEX6bp vector containing GST-3C tag by BamH1-Not1. USP1 amino acids 1–60 N-terminal was amplified with BamH1 at either end for ligation into a pGex6bp1 vector containing rat USP2 catalytic domain (GST-3C-USP1^1^-60-rUSP2 (271–618)). A mutant Ube2T with E54R, P93G, P94G, 1–152 (Ube2T^V3^)^33^. FANCL^ΔELF^ (109–375) was cloned into a pET-SUMO (Invitrogen) vector using restriction free cloning. Tagless Ubiquitin, His-TEV Ubiquitin and His-TEV-PCNA are previously described^22,44^. RNF4 (RING-RING domain fusion^34^) was a kind gift from Ron Hay (University of Dundee). To introduce point mutations or truncations, PCR mutagenesis was performed using KOD Hot Start DNA Polymerase following manufacturer’s instructions.

### Protein expression and purification

Baculovirus were generated using the EMBacY MultiBac system (Geneva biotech) and USP1, UAF1, FANCD2 and FANCI proteins were expressed for ∼72h in baculovirus infected sf21 cells. FANCL^ΔELF^, PCNA, ubiquitin constructs, Ube2T^V3^ and RNF4^RING^-RING were expressed and purified in BL21 *E.coli* as previously described^34,44^. USP1^Δ1Δ2^ and USP2 were expressed using BL21 *E.coli*, grown to OD600: 0.6 and induced with 0.1 µM Isopropyl-Beta-d-Thiogalactopyranoside (IPTG) for 18h at 16°C. Fluorescently labelled ubiquitin was made as described previously^42^.

All steps of purifications were performed at 4°C and completed within 24–36 hours of lysis. Cell pellets were routinely suspended in lysis buffer (50 mM Tris pH 8.0, 150 mM NaCl, 5% glycerol, 10 mM imidazole, 10 mM 2-Mercaptoethanol with freshly added MgCl2 (2 mM), protease inhibitor (EDTA-free) tablets and benzonase. For FANCD2, FANCI and FANCL^Δ^ELF and lysis buffer contained 400 mM NaCl. Sf21 cells were lysed by homogeniser followed by sonication at 40% amplitude, 10s on 10s off for 12 cycles. Suspended *E.coli* cells were sonicated at 80% amp, 20s on 40s off, 12 cycles. All lysates were clarified at 40,000 x g for 45 min and filtered (0.45 µM). Proteins were bound to respective resins (either Ni-NTA for His tags or GSH for GST tags) and washed extensively with lysis buffer with 500 mM NaCl. GST tagged proteins were eluted by incubation with GST-precision protease or lysis buffer with 10 mM GSH. His-Smt3 tagged proteins were eluted by on column cleavage using His-ULP1 protease. His-TEV or His -3C tagged proteins were eluted by lysis buffer with 250 mM imidazole and lower NaCl for ion exchange chromatography (typically 100 mM NaCl). To remove tags, proteins were incubated with respective proteases (His-TEV protease or GST-precision protease) overnight and tagged proteases were removed by binding to respective resin. Tags were not removed from FANCD2, FANCI, USP2 or GST-Ubiquitin. Anion exchange for USP1, UAF1, FANCD2, FANCI, GST-USP2, GST-Ub and PCNA were all performed using a HPQ (1 ml or 5 ml) column and eluted with a linear gradient (50 mM Tris pH 8.0, 100–1000 mM NaCl, 5% glycerol and 10 mM 2-Mercaptoethanol). *E.coli* expressed USP1 was not purified by anion exchange.

Proteins were further purified using size exclusion chromatography (SEC). FANCD2 and FANCI were fractionated using an S6 Increase (10/300) column in 20 mM Tris pH 8.0, 400 mM NaCl, 5% glycerol, 5 mM DTT. USP1, UAF1 and E3s were fractionated using an SD200 Increase (10/300) in 20 mM Tris pH 8.0, 150 mM NaCl, 5% glycerol with 5 mM DTT. All ubiquitin constructs and E2s were fractionated using an SD75 (10/300) in 20 mM Tris pH 8.0, 150 mM NaCl, 5% glycerol, 0.5 mM Tris 2-carboxyethyl-phosphine (TCEP). Proteins were concentrated to ∼5 mg/ml for FANCD2/FANCI, ∼4 mg/ml for FANCL^Δ^ELF and ∼10 mg/ml for all other proteins before flash freezing in liquid nitrogen and storage at -80°C.

### Analytical Gel filtration of USP1-UAF1 complexes

Recombinant USP1 and UAF1 alone or together were incubated at 15 µM for 15 min on ice before fractionation on SD200 Increase (10/300) in 20 mM Tris pH 8.0, 150 mM NaCl, 5% glycerol and 5 mM DTT. Samples (10 µl) were mixed with SDS-page loading buffer (10 µl) and analysed via SDS-page (9%) and instant blue coomassie staining (Expedeon). All USP1 constructs co-eluted with UAF1.

### Ubiquitin-propargylamine (Ub-prg)

Ub-intein-CBD in BL21 *E.coli* was grown to OD600: 0.6, induced with 0.5 µM IPTG and expressed for 24 h at 16°C. Cells were harvested and suspended in 50 mM Na2HPO4 pH 7.2, 200 mM NaCl, 1 mM EDTA before lysis and clarification. Lysates were bound to chitin-resin and washed extensively in 50 mM Na2HPO4 pH 7.2, 200 mM NaCl. Resin was washed with wash buffer (20 mM Na2HPO4 pH6.0, 200 mM NaCl, 0.1 mM EDTA) before incubation overnight at 4°C with wash buffer with 100 mM MESNa. Elute was collected and a second elution was performed in wash buffer plus 100 mM MESNa. The Ubiquitin-MESNa was concentrated to ∼5 ml and buffer exchange into 50 mM HEPES pH 8.0. Ubiquitin-MESNA was either stored at -80°C or directly reacted with 0.25 M propargylamine (prg) for 4 h at 20°C with mild shaking in the dark. Ub-prg was finally purified using SD75 (16/60) in 50 mM HEPES pH 8.0, 150 mM NaCl and flash frozen for storage at -80°C.

### Reacting Ub-prg with DUBs

To crosslink DUBs with Ub-prg, DUBs (2 µM) in 20 µl were incubated in DUB buffer (50 mM Tris pH 7.5, 120 mM NaCl, 10 mM DTT) with and without Ub-prg (6 µM) for 10 min at room temperature. Reactions were stopped by addition SDS-page loading buffer (Invitrogen) and visualised using 4–12% Bis-Tris SDS-PAGE gel (Invitrogen) run with NuPAGE™ MOPS SDS Running Buffer (Thermo Scientific) and coomassie staining a slower migrating band indicates reaction with Ub-prg.

### Thermofluor assays

Thermofluor experiments were carried out using a CFX96 real-time PCR detection system (Bio-Rad). USP1 (40 µl) at 0.05–0.2 mg/ml in 50 mM Tris pH8.0, 120 mM NaCl, 10 mM DTT and 5x SYPRO orange was prepared in 96-well plates. The samples were heated from 25°C to 95°C with increments of 1°C/minute, and fluorescence was measured at each interval. Data was analysed as previously described^45^ and a mean T_m_°C was calculated from three different USP1 concentrations which were performed in triplicate.

### Fluorescent polarisation (FP) assays with Ubiquitin-TAMRA assays

USP1-UAF1 were assembled by diluting USP1 variants (10 µM) and UAF1 (10 µM) in DUB buffer for 10 min at room temperature. For FP assays, USP1-UAF1 were diluted and incubated in DUB buffer with 0.1 mg/ml ovalbumin at 2X concentration indicated in reaction. Typically, reactions with Ub-KG^TAMRA^ (UbiQ) were started by adding and mixing 10 µl of 2X DUB (20 nM) to 10 µl 2X Ub-KG^TAMRA^(600 nM) to give a final concentration of 300 nM substrate with 10 nM DUB with a volume of 20 µl. Reactions were monitored at 25°C for 1h by measuring FP at 2 min intervals in 384-well round bottom corning black plates with a PHERAstar FSX. FP values for each well were fitted using ‘one phase decay’ model in Prism (GraphPad). A pMax was monitored using Ub-KG^TAMRA^ without DUB and pMin using KG^TAMRA^.

### Purifying monoubiquitinated proteins

Purification of PCNA monoubiquitinated at K164 (PCNA-Ub) was performed as previously described^25^. Monoubiquitinated FANCD2 at K561 and FANCI at K523 was purified using GST-Ub which leaves a ‘GPLGS’ over hang at the N-terminus of ubiquitin following precision protease cleavage. Typically, purified FANCD2 or FANCI (after anion exchange and relatively fresh protein ∼4h from cell lysis) at 4 µM was monoubiquitinated by equimolar Ube2T^V3^, FANCL^Δ^ELF, 50 nM E1 and 8 µM GST-3C-ubiquitin in E3 buffer (50 mM Tris pH 8.5, 150 mM NaCl, 5% glycerol, 1 mM TCEP, 2.5 mM MgCl2 and 2.5 mM ATP) for 30 min at room temperature. *Xl* FANCD2 ubiquitination was performed for 10 min rather than 30 min. Reactions were arrested using apyrase followed by anion exchange chromatography (HPQ 1ml). Importantly, prolonged incubation of FANCD2 in low NaCl concentrations (below ∼200 mM NaCl) will lead to large losses in protein. Since, ubiquitinated substrates eluted together with non-ubiquitinated substrate from anion exchange, GST-Ub-Substrates were bound to GSH resin at 4°C before extensive washes using 50 mM Tris pH 8.0, 400 mM NaCl, 5% Glycerol and 2 mM DTT. Ubiquitinated proteins were eluted using GST-precision protease by incubation with GSH-resin at 4°C. Finally, ubiquitinated-Substrates were purified via gel filtration S6 (10/300) in 20 mM Tris pH 8.0, 400 mM NaCl, 5% glycerol, 5 mM DTT before concentrating to ∼5 mg/ml and flash freezing in liquid nitrogen for storage at -80°C. To determine difference between ubiquitinated and non-ubiquitinated FANCD2 or FANCI, proteins were visualised on Bis-Tris 4–12% gradient gels run in MOPs running buffer, gels were ‘overrun’ so the 20 kDa reference band in the precision plus pre-stained protein ladder (Bio-Rad) was at the bottom of the SDS-PAGE gel. To purify FANCD2-Ub^800^, FANCD2 was monoubiquitinated with Ubiquitin^800^instead of GST-Ub, and subsequently purified by anion exchange and gel filtration as above.

### Mass spectrometry

FANCD2, FANCI, FANCD2-Ub and FANCI-Ub (10 µg) were denatured in 7M Urea, 100 mM Tris pH 8.0 buffer followed by treatment with 5 mM TCEP (10 min), 5 mM Idoacetamide (20 min) and 5 mM DTT (20 min). Samples were diluted 1:10 in 100 mM Tris pH 8.0 and trypsin was added to 1:50 w/w with incubation at 37°C o/n, finally 1% TFA was added. The peptides were subjected to mass spectrometric analysis performed by LC-MS-MS on a linear ion trap-orbitrap hybrid mass spectrometer (Orbitrap-VelosPro, Thermo) coupled to a U3000 RSLC HPLC (Thermo). Data files were analysed by Proteome Discoverer 2.0(www.ThermoScientific.com), using Mascot 2.4.1 protein identification software (www.matrixscience.com) by searching against an in house protein database.

### Deubiquitination reactions using recombinant substrates

USP1-UAF1 complexes are assembled as above and used to make 2X DUB stocks at 200 nM or as indicated in figure. Ubiquitinated-substrates were made as 2X stocks: purified *hs*FANCD2-Ub (2 µM), *hs*FANCI-Ub (2 µM), *hs*PCNA-Ub (6 µM), *x*/FANCD2-Ub (2 µM) and diubiquitin chains (20 µM) were incubated in DUB buffer on ice for 10 min. To initiate hydrolysis, DUBs (200 nM) and substrates (2 or 6 µM) were incubated in a 1:1 ratio (typically 5 µl: 5 µl) for 30 min at room temperature or an indicated time (typically 30 min). Reactions were terminated using SDS-loading buffer (Invitrogen). To analyse deubiquitination, ∼ 300 ng of *hs*FANCD2-Ub or *hs*FANCI-Ub, ∼600 ng of *hs*PCNA-Ub or 800 ng of diubiquitin was separated on 4–12% Bis-Tris SDS-PAGE gels and coomassie staining. Deubiquitination was determined by the loss of ubiquitinated protein or the emergence of non-ubiquitinated substrate and monoubiquitin.

### Ubiquitination reactions

FANCD2 at K561 and FANCI at K523 was ubiquitinated at room temperature in E3 reaction buffer using Ube2T^V3^ and FANCL^Δ^ELF. For small scale reactions (10–100 µl) proteins concentrations were typically 2 µM substrate/E2/E3, 4 µM Ubiquitin and 50 nM E1. To Ubiquitinate FANCD2 at alternative lysines to K561 (i.e. FANCD2-KX-Ub), FANCD2 K561R was ubiquitinated using 50 nM E1, 4 µM Ubiquitin^800^, UbcH5c (S22R) 0.5 µM and 0.2 µM RNF4^RR^ in E3 reaction buffer at room temperature. Reactions were stopped by addition of 3X SDS-PAGE loading buffer or, if used for subsequent DUB reactions, was apyrase treated for 5 min on ice. Ubiquitination was analysed via 4–12% Bis-Tris SDS-PAGE, Odyssey licor CLx with 800 nm channel for Ubiquitin^800^ and coomassie staining.

### DUB-Step reactions

DUB-step reactions followed 3 steps: 1) ubiquitination (see- *Ubiquitination reactions*), 2) arrest with apyrase to stop ubiquitination and 3) deubiquitination. Briefly, FANCD2 (2 µM) ubiquitinated on K561 or KX was immediately diluted 5 µl: 5 µl with DUBs (200 nM) diluted in DUB buffer, giving a final concentration of DUB at 100 nM and FANCD2 at 1 µM. DUB-steps were left at room temperature for indicated amount of times (usually 30 min) and 300 ng of FANCD2-Ub^800^ was visualised on SDS-PAGE (4–12% Bis-Tris) with 800 nM channel followed by coomassie staining. The percentage of residual FANCD2-Ub^800^ was calculated as a percentage of the input and plotted for each DUB.

### Pull down assays

N-terminal strep-tagged-UAF1 was complexed with USP1^Δ1Δ2^ or USP1^Δ^NΔ1Δ2. Subsequently, 10 µg of USP1-UAF1 was bound to 10 µl MagStrep-avidin beads equilibrated in pull down buffer (50 mM Tris pH 7.5, 120 mM NaCl, 5% glycerol, 1 mM DTT, 0.01% Triston X-100 and 0.01% ovalbumin) for 1 h on ice with gentle agitation. Unbound USP1-UAF1 complexes were washed from beads with 3X 60 µl washes. USP1-UAF1 beads were then divided into separate tubes (10 µl beads + 10 µl buffer) and 10 µg of recombinant substrate was added and incubated for 1 h on ice. Supernatant was removed and beads were washed 3 x with 40 µl buffer. Protein was eluted by suspending beads in 20 µl pull down buffer and 10 µl 3X SDS-PAGE loading buffer and boiling for 2 min. Samples were run on 4–12% Bis-Tris SDS-PAGE with 10 µl of sample. Beads without USP1-UAF1 were also incubated with substrate to control for non-specific binding to beads.

### Western blots and antibodies

SDS-PAGE separated proteins were transferred to nitrocellulose membranes and blocked with 5% milk TBS-T (0.05% tween) before incubation with 1:1000 anti-FANCD2 (Ab2187, Abcam), 1: 5000 anti-V5 (66007.1- Ig, ProteinTech) or 1: 2000 anti-PCNA (Ab29, Abcam) overnight at 4 °C. Membranes were washed 3X with TBST before incubation with secondary antibodies for 1h at room temperature and washes with TBS-T. Results were visualised on licor using 800 nm Channel.

## Ackowledgements

We thank Axel Knebel (University of Dundee) for His-Ube1. We thank Robert Gourlay from the MRC PPU Proteomics Facility (university of Dundee) for mass spectrometry. This work was supported by the Medical Research Council (MRC grant number MC_UU_12016/12); the EMBO Young Investigator Programme to H.W.; the European Research Council (ERC-2015-CoG-681582 ICLUb consolidator grant to H.W.

## Author Contributions

CA purifed proteins, designed and performed all experiments. VKC purfied proteins and designed experiments. CA, RT and VKC cloned several constructs. CA and HW analysed data and wrote the manuscript.

